# Thyroid hormone regulates distinct paths to maturation in pigment cell lineages

**DOI:** 10.1101/527341

**Authors:** Lauren M. Saunders, Abhishek K. Mishra, Andrew J. Aman, Victor M. Lewis, Matthew B. Toomey, Jonathan S. Packer, Xiaojie Qiu, José L. McFaline-Figueroa, Joseph C. Corbo, Cole Trapnell, David M. Parichy

## Abstract

Circulating endocrine factors are critical for orchestrating complex developmental processes during the generation of adult form. One such factor, thyroid hormone, regulates diverse cellular events during post-embryonic development and can drive disparate morphological outcomes through mechanisms that remain essentially unknown. We sought to define how thyroid hormone elicits opposite responses in the abundance of two pigment cell classes during development of the zebrafish adult pigment pattern. By profiling individual transcriptomes from thousands of neural crest derived cells, including pigment cells, we reconstructed developmental trajectories and identified lineage-specific changes in response to thyroid hormone. Contrary to our initial hypotheses for how TH differentially affects two related pigment cell lineages, we find instead that TH promotes the maturation of both melanophores and xanthophores in distinct ways, promoting cellular senescence and carotenoid accumulation, respectively. Our findings show that thyroid hormone and its receptors regulate distinct events of cellular maturation across lineages, and illustrate how a single, global factor integrates seemingly divergent morphogenetic outcomes across developmental time.

## Introduction

Mechanisms that synchronize developmental signals and integrate them across cell types and organ systems remain poorly defined but are fundamentally important to both development and evolution of adult form (Atchley and Hall, 1991; Ebisuya and Briscoe, 2018). A powerful system for elucidating how organisms coordinate fate specification and differentiation with morphogenesis is the array of cell types that arise from embryonic neural crest (NC), a key innovation of vertebrates (Gans and Northcutt, 1983). NC cells disperse throughout the body, contributing peripheral neurons and glia, osteoblasts and chondrocytes, pigment cells and other derivatives. Differences in the patterning of these cells underlie much of vertebrate diversification.

Thyroid hormone (TH) coordinates post-embryonic development of NC and other derivatives through mechanisms that are incompletely characterized (Brown and Cai, 2007; Shi, 1999). During the abrupt metamorphosis of amphibians, TH drives outcomes as disparate as tail resorption and limb outgrowth (Shi, 1999). In the more protracted post-embryonic development of zebrafish—which has similarities to fetal and neonatal development of mammals (Parichy et al., 2009)—TH coordinates modifications to several traits including pigmentation. Remarkably, TH has seemingly opposite effects on two classes of NC-derived pigment cells, curtailing the population of black melanophores yet promoting development of yellow/orange xanthophores; fish lacking TH have about twice the normal number of melanophores and lack visible xanthophores (**Figure 1A**) (McMenamin et al., 2014).

**Figure 1.**
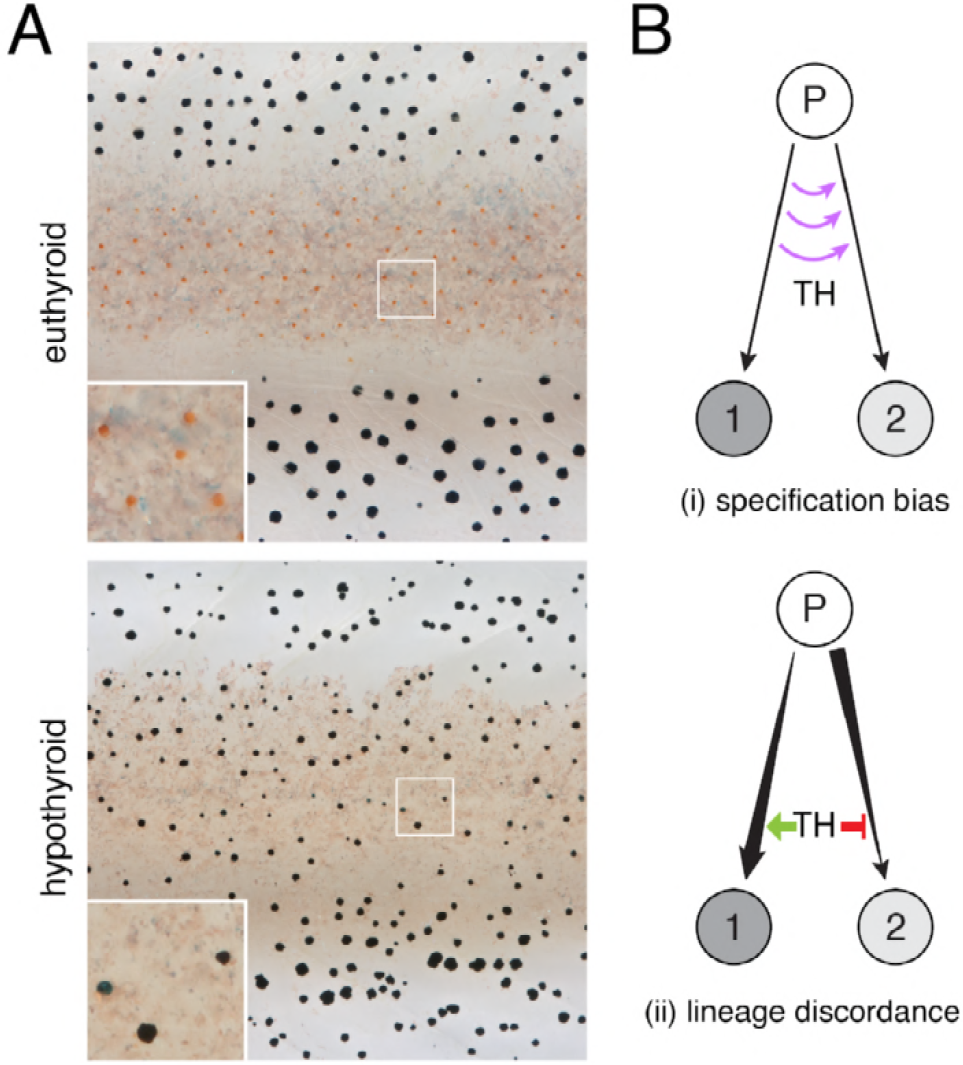
TH dependent phenotypes and models for TH action. **(A)** Euthyroid and hypothyroid zebrafish [stage 10 SSL (Parichy et al., 2009); ~21 d post fertilization, dpf]. Insets, yellow/orange xanthophores of euthyroid fish and absence of these cells in hypothyroid fish. **(B)** Models for TH effects on alternative cell types derived from a common progenitor (P), by regulating cell fate specification (i) or lineage-specific amplification and restraint of committed cell-types, through differential effects on morphogenesis and differentiation (ii).

We asked how a single endocrine factor can have such different effects on cells sharing a common embryonic origin. Using transcriptomic analyses of individual cell states, we comprehensively define the context for TH activities by identifying populations and subpopulations of adult NC derivatives. We then assess the consequences of TH status for lineage maturation across pigment cell classes. Our analyses show that TH drives maturation of cells committed to melanophore and xanthophore fates through different mechanisms, involving cellular senescence and carotenoid-dependent repigmentation, respectively. These mechanisms reflect different developmental histories of these NC sublineages, and yield different cell-type abundances when TH is absent. Our findings provide new insights into post-embryonic NC lineages, contribute new resources for studying adult pigment cells and other NC-derived cell types, and illustrate how a circulating endocrine factor influences local cell behaviors to coordinate adult trait development.

## Results

### Post-embryonic NC-derived subpopulations revealed by single cell RNA-sequencing

To explain the pigment cell imbalance of hypothyroid fish, we envisaged two models for TH activity during normal development (**Figure 1B**): (i) TH influences specification, directing multipotent cells away from one fate and towards the other; or (ii) TH has discordant effects on different lineages, driving the selective amplification of cells already committed to one fate while simultaneously restraining amplification of cells committed to the other fate.

To evaluate the contributions of these two models to the imbalance of pigment cells, we sought to capture the range of intermediate states through which these cells transit during normal and hypothyroid development. Accordingly, we sequenced transcriptomes of thousands of individual NC-derived cells isolated from trunks of euthyroid and hypothyroid fish (**Figure 2—figure supplements 1** **and** **2**). Dimensionality reduction (Becht et al., 2018; Cao et al., 2018) followed by unsupervised clustering identified melanophores, xanthophores and a third class of NC-derived pigment cells, iridescent iridophores (**Figure 2A** **and B**; **Supplementary HTML**). A cluster likely corresponding to multipotent pigment cell progenitors (Kelsh et al., 2017; Singh et al., 2016) was marked by genes encoding pigment cell transcription factors, mesenchymal NC markers, and factors associated with proliferation and migration but not pigment synthesis (**Figure 2—figure supplement 3**; **Table 1**). Also present were neurons, Schwann cells and other glia, and presumptive progenitors not described previously. Clusters exhibited distinct expression of genes encoding ligands and receptors, cell adhesion molecules, and products likely to have diverged in function after genome duplication (**Figure 2—figure supplement 4**). To learn if NC-derived cells employ similar gene expression programs across life cycle phases, we compared cells isolated from embryonic–early larval (“EL”) and middle larval–juvenile (“adult”) stages, which revealed broad overlap of transcriptional profiles (**Figure 2—figure supplement 5**). Overall, our survey captured numerous NC-derived cell types, including abundant pigment cells and progenitors, and revealed substantial variation in gene expression programs among them.

**Figure 2.**
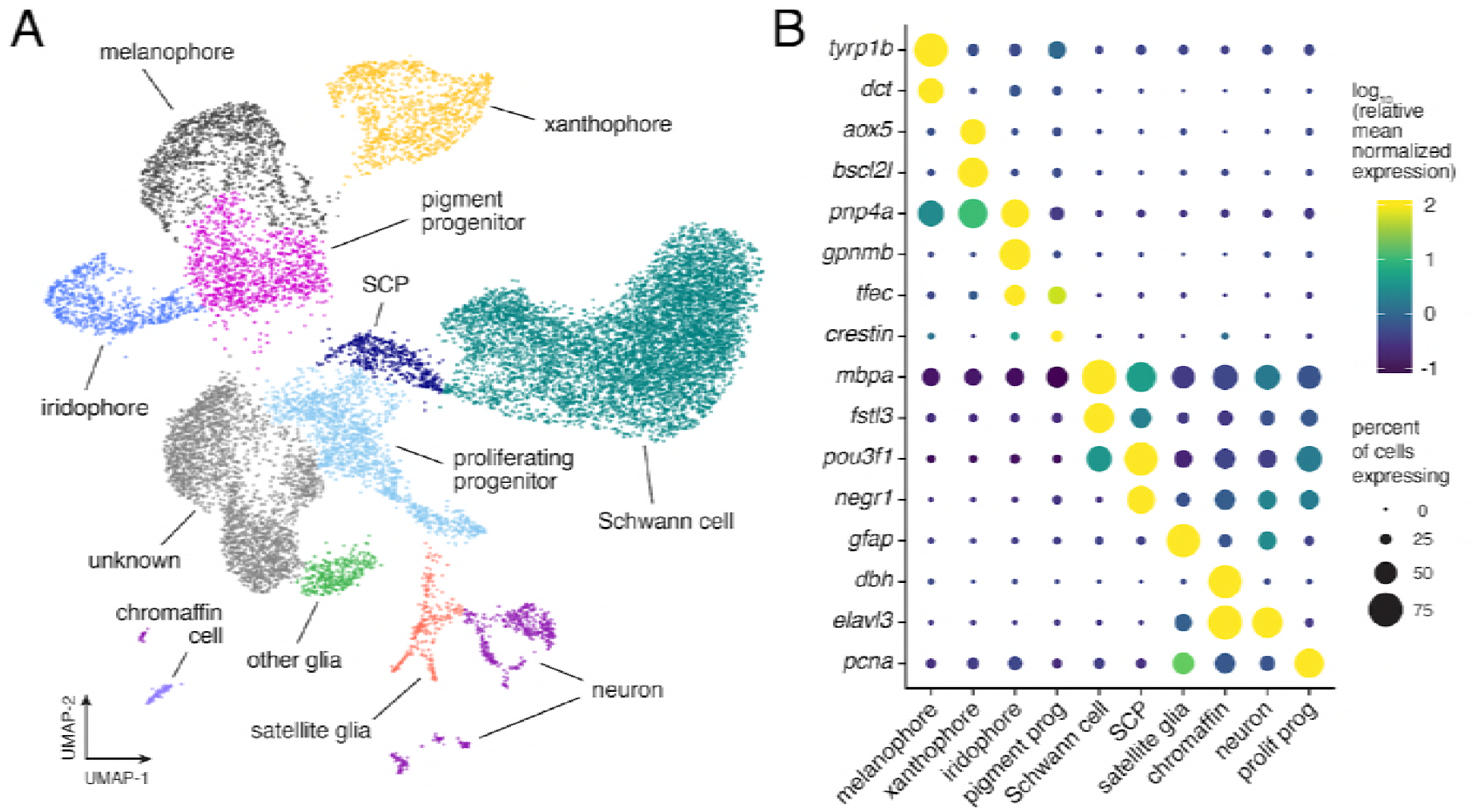
Single cell transcriptomic identification of post-embryonic NC-derived cell types. **(A)** Cell-type assignments for clusters of cells (*n*=16,150) from euthyroid and hypothyroid fish. Cell types known to be of non-NC derivation were removed from analysis. **(B)** Known cell-type marker genes and new candidate markers (for cluster-specific genes, see **Table 1**).

### Pigment cell sub-classes and gene expression dynamics across differentiation

To understand the gene expression context in which TH impacts each pigment cell type, we compared pigment cells and progenitors, the lineages of which have been described (Budi et al., 2011; Eom et al., 2015; Mahalwar et al., 2014; McMenamin et al., 2014; Singh et al., 2016) (**Figure 3A**). These analyses revealed subsets of melanophores and xanthophores (**Figure 3B; Figure 3—figure supplement 1**), consistent with distinct morphogenetic and differentiative behaviors (Eom et al., 2015; Parichy et al., 2000b; Parichy and Spiewak, 2015), new markers of xanthophore and iridophore lineages (**Figure 3—figure supplements 2** **and** **3**), and cell-type specific expression of some previously identified markers [e.g., *tyrp1b*, *aox5*, *tfec* (Lister et al., 2011; McMenamin et al., 2014)] (**Figure 3C**). Expression of other genes was broader than might be expected from mutational or other analyses (**Figure 3—figure supplement 4**); e.g., *mitfa*, encoding a transcription factor required for melanophore fate specification (Lister et al., 1999) was expressed in melanophores and progenitors, but also xanthophores (**Figure 3C**).

**Figure 3.**
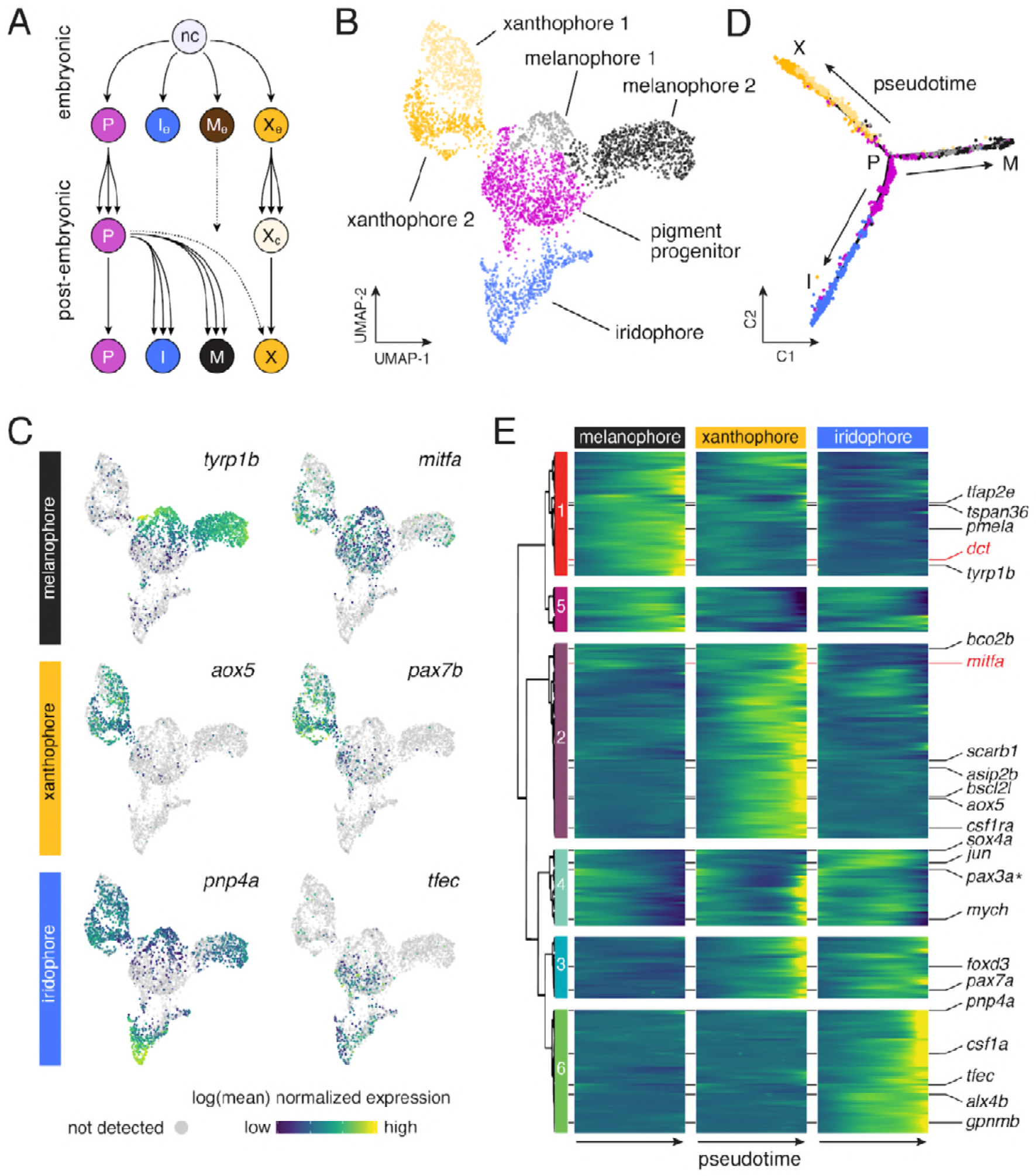
Pigment cell subpopulations and dynamics of gene expression across pigment cell lineages. **(A)** Established lineage relationships of embryonic (e) and post-embryonic pigment cells. Multipotent pigment cell progenitors (P) in the peripheral nervous system generate adult iridophores (I), melanophores (M) and some xanthophores (X). A few embryonic melanophores (M_e_) persist whereas embryonic xanthophores (X_e_) proliferate and lose their pigment to enter a cryptic phase (X_c_), and then reacquire pigment late in pattern formation to form most adult xanthophores(Patterson et al., 2014). **(B)** Sub-clusters of melanophores and xanthophores with distinct gene expression signatures. **(C)** Pigment cell clusters defined by markers for each cell-type (Parichy and Spiewak, 2015). **(D-E)** Pseudotemporal ordering (D) and BEAM (E) revealed dynamics of gene expression over pseudotime for each pigment cell branch (*q*<6.0E-11 for all genes; except *pax3a* (starred, *q*=0.03), expressed as anticipated during early pseudotime in each branch.

To characterize transcriptional dynamics through lineage maturation, we pseudotemporally ordered cells (Qiu et al., 2017a, 2017b; Trapnell et al., 2014), yielding a differentiation trajectory with each pigment cell type arising from a common progenitor (**Figure 3D**). This topology differed from lineage relationships (**Figure 3A**), but was consistent with similarity of EL and adult gene expression programs (**Figure 2—figure supplement 5D**). Branch expression analysis modeling (BEAM) (Qiu et al., 2017a) confirmed that genes known to function in specification (e.g., *mitfa* in melanophores) were expressed early in pseudotime whereas genes associated with differentiation [e.g., *dct*, encoding a melanin synthesis enzyme (Kelsh et al., 2000b)] were expressed late (**Figure 3E**; **Figure 3—figure supplement 5A**). These analyses revealed dynamics of hundreds of genes likely identifying discrete processes in lineage-specific maturation (**Table 2**) as well as broader trends. For example, transcripts per cell declined in melanophores but not iridophores, consistent with an expectation of reduced RNA abundance as melanophores—but not iridophores—exit the cell cycle with maturation (**Figure 3—figure supplement 5B**) (Budi et al., 2011; Darzynkiewicz et al., 1980; McMenamin et al., 2014; Spiewak et al., 2018).

### TH-independence of pigment cell fate specification and absence of lineage-specific restraints on developmental progress

Resolution of pigment cell states through their development allowed us to test if TH functions in fate specification (**Figure 1B-i**). If so, excess melanophores and missing xanthophores of hypothyroid fish should reflect biases on specification of multipotent progenitors or transdifferentiation of initially specified cells. Such alterations should be evident in reduced-dimension transcriptomic space as strong skew in the apportionment of cells between branches or abnormal paths in the cellular trajectory, respectively. Yet, euthyroid and hypothyroid trajectories were topologically equivalent. Moreover, pigment cell progenitors were not depleted in hypothyroid fish as might occur were these cells being allocated inappropriately as melanophores (**Figure 4A–D**).

**Figure 4.**
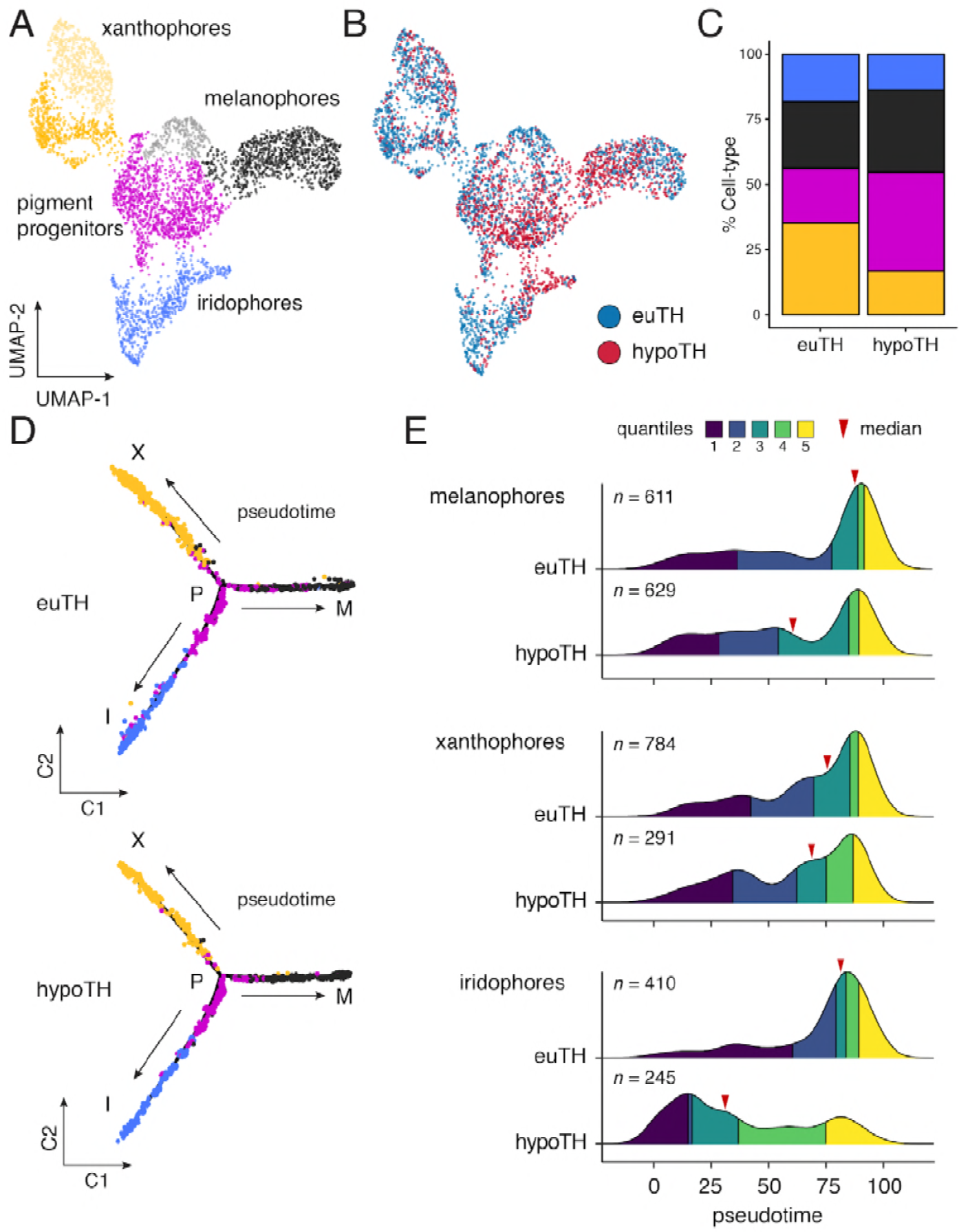
Thyroid hormone biased pigment cell lineages towards later steps of pseudotime. **(A)** UMAP dimensionality reduction of euthyroid and hypothyroid pigment cells and pigment progenitors. Sub-clustering reveals two xanthophore and two melanophore clusters (indicated by different shades of yellow and grey, respectively). **(B)** Pigment cells colored by TH status. Euthyroid and hypothyroid cells generally intermix with some biases apparent within melanophore and iridophore clusters. **(C)** Percentages of each pigment cell class by TH-status. Colors are consistent with other pigment cell plots. Of cells captured, a higher proportion of pigment cells from euthyroid fish were xanthophores and iridophores compared to those from hypothyroid fish. **(D)** Trajectories for euthyroid and hypothyroid pigment cells (plotted together, faceted by condition). Broad differences in trajectory topologies were not apparent between the two conditions. **(E)** Distributions of each pigment cell-type across pseudotime by condition. For each trajectory branch (mel, xan, irid) hypoTH cells were biased towards early pseudotime (Wilcoxon signed-rank tests, mel: Z=−6.54, *P*<0.0001, xan: Z=−4.54, *P*<0.0001, irid: Z=−13.55, *P*<0.0001). Median is indicated by red arrowhead and different colors demarcate quartiles over pseudotime.

Through a second model—lineage discordance—TH could have opposite effects on cells already committed to particular fates, selectively amplifying one cell type while simultaneously repressing amplification of the other (**Figure 1B-ii**). For example, TH could promote differentiation of xanthoblasts to xanthophores, but prevent differentiation of melanoblasts to melanophores. Alternatively, TH could be a survival factor in the xanthophore lineage but a pruning factor in the melanophore lineage. Terminal phenotypes of both hypothyroid and hyperthyroid mutant fish are consistent with such effects (McMenamin et al., 2014). If TH has discordant effects between lineages, we predicted that hypothyroid fish should exhibit a strong depletion of xanthophores from the end of their branch of the trajectory, whereas melanophores should be strongly over-represented near the tip of their branch. Yet, empirical distributions of pigment cell states in hypothyroid fish were all biased towards earlier steps in pseudotime, sometimes severely (**Figure 4E**). Indeed, prior analyses showed that addition of exogenous TH to hypothyroid cells *ex vivo* can promote differentiation of unpigmented melanoblasts to melanophores (McMenamin et al., 2014), contrary to the idea that TH specifically blocks melanophore development. Together these findings allow us to reject a model in which TH regulation of pigment cell abundance in the adult fish depends on discordant effects across lineages.

### TH promotes a melanophore maturation program and drives cells to a senescent state

Having rejected both of our initial models (**Figure 1B**), we considered a third possibility, that TH promotes the maturation of both lineages, but in distinct ways. For melanophores, inspection of transcriptomic states and cellular phenotypes strongly supported a role for TH in promoting maturation of this lineage. Genes expressed during terminal differentiation of melanophores from euthyroid fish were expressed at lower levels in melanophores of hypothyroid fish, suggesting an impediment to maturation in the absence of TH (**Figure 5A**). Cytologically, mature melanophores of euthyroid fish are frequently binucleate—a condition associated with increased survival and larger size (Orr-Weaver, 2015; Usui et al., 2018)—whereas we found that melanophores of hypothyroid fish were frequently mononucleate, suggesting they had failed to complete differentiation (**Figure 5B** **and C**). Given these findings and an earlier observation that melanophores of euthyroid fish fail to divide whereas those of hypothyroid fish continue to do so (McMenamin et al., 2014), we hypothesized that TH promotes a terminally differentiated state of cellular senescence. Consistent with this idea, mature melanophores often exhibited senescence-associated β-galactosidase activity, and melanophores isolated from euthyroid fish had significantly greater senescence-associated lysosomal content (Kurz et al., 2000) than melanophores from hypothyroid fish (**Figure 5D** **and E**; **Figure 5—figure supplement 1**). These results indicate that TH drives melanophores to a senescent state. Excess melanophores of hypothyroid fish thus reflect an immature, amplifying phenotype retained even after melanization.

**Figure 5.**
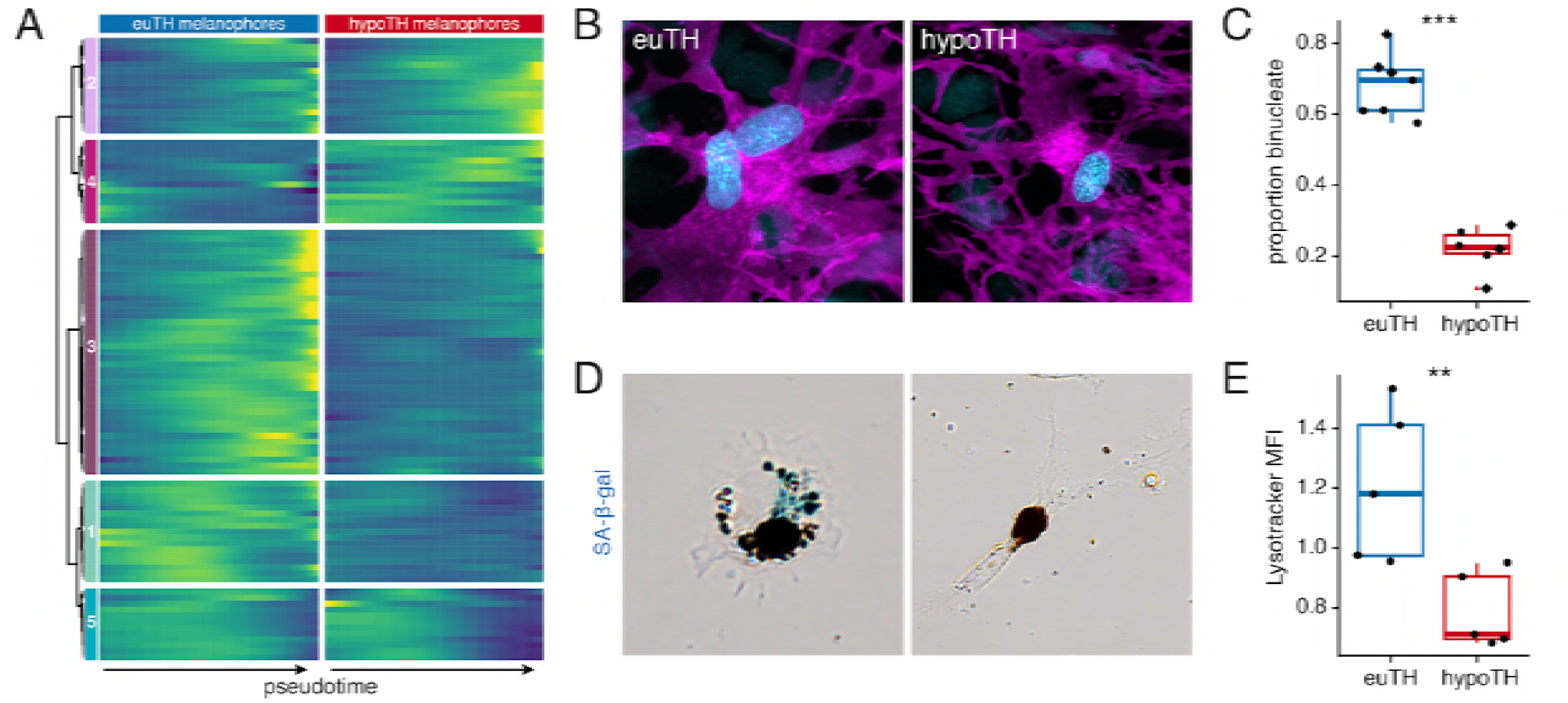
TH promoted melanophore maturation by measures of transcriptomic state and cellular phenotype. **(A)** Gene expression differences between melanophores over pseudotime by TH-status (*q*<1E-7, genes expressed in >10% of melanophores). Heatmap is hierarchically clustered by row (method, Ward D2). The largest cluster (#3) contains 41% of the genes and represents loci expressed late in pseudotime of euthyroid melanophores but downregulated in hypothyroid melanophores, identifying novel candidate genes for roles in melanophore maturation (see **Table 6**, in which published melanophore-related genes are highlighted) (Baxter et al., 2018). **(B)** Fully differentiated melanophores of zebrafish are often binucleate (Usui et al., 2018) as, shown *in vivo* for a stripe melanophore isolated from a euthyroid fish (12 SSL). By contrast, melanophores of hypothyroid fish at equivalent stages were often mononucleate. Magenta, membrane labeling of melanophores by *tyrp1b:palm-mCherry*. Blue, nuclei revealed by *tuba8l3:nEosFP*. **(C)** Hypothyroid fish had proportionally fewer binucleate melanophores than euthyroid fish (*χ*^2^=230.3, d.f.=1, *P*<0.0001) after controlling for a higher incidence of binucleation in developmentally more advanced fish overall (11.5–13 SSL; *χ*^2^=5.5, d.f.=1, *P*<0.05). Individual points indicate proportions of binucleate melanophores observed in dorsal stripes (circles) and ventral stripes (diamonds), which did not differ significantly (*P*=0.8; sample sizes: *n*=383 melanophores in 4 euthyroid fish, *n*=706 melanophores in 3 hypothyroid fish). **(D)** Melanophores plated *ex vivo* from fish at terminal stages of stripe development had diverse morphologies and some exhibited senescence-associated β-galactosidase staining (blue precipitate, left panel). Xanthophores plated *ex vivo* did not exhibit senescence-associated β-galactosidase staining. **(E)** Cellular senescence assayed quantitatively by lysosomal content (Kurz et al., 2000; Lee et al., 2006) revealed significantly greater normalized mean fluorescence intensity (MFI) of Lysotracker dye in melanophores isolated from euthyroid fish as compared to hypothyroid fish (euthyroid, 20,423 melanophores; hypothyroid, 87,252 melanophores; *P*<0.01, Wilcoxon).

### TH promotes carotenoid-dependent xanthophore re-pigmentation during adult development

We next examined TH functions specific to the xanthophore lineage. Most adult xanthophores develop directly from EL xanthophores that lose their pigment and then reacquire it late in adult pattern formation (**Figure 3A**) (McMenamin et al., 2014). Because xanthophores of hypothyroid fish persist, albeit in a cryptic state (McMenamin et al., 2014), we predicted that TH effects should be less pervasive in these cells than in melanophores that develop *de novo* from a post-embryonic progenitor. Indeed, fewer genes were expressed differentially between TH backgrounds in xanthophore than melanophore lineages (3.6% vs. 9%; **Figure 6A**). Prominent among these were several loci implicated in, or plausibly associated with, yellow/orange carotenoid coloration (**Figure 6B** **and C;** **Figure 6—figure supplement 1**) (Toews et al., 2017). These differences in gene expression suggested a carotenoid pigmentation deficiency that we confirmed by HPLC, histology, and transmission electron microscopy (**Figure 6D; Figure 6—figure supplement 2**). Among carotenoid genes, *scavenger receptor B1* (*scarb1*) encodes a high density lipoprotein receptor essential for carotenoid accumulation in birds and invertebrates (Kiefer et al., 2002; Toomey et al., 2017) and we found it to be required in zebrafish for carotenoid deposition, though not cell persistence (**Figure 6—figure supplement 3A** **and B**). *scarb1* was expressed more highly in xanthophores of euthyroid than hypothyroid fish (*q*=1.1E-10) (**Figure 6B** **and E**; **Figure 5—figure supplement 1**) and exogenous TH was sufficient to rescue both expression and yellow/orange coloration (**Figure 6F; Figure 6—figure supplement 3C**). Together these findings demonstrate an essential role for TH in carotenoid pigmentation and suggest that concerted TH modulation of a suite of carotenoid pathway genes is required for cryptic xanthophores to re-pigment during adult pattern formation.

**Figure 6.**
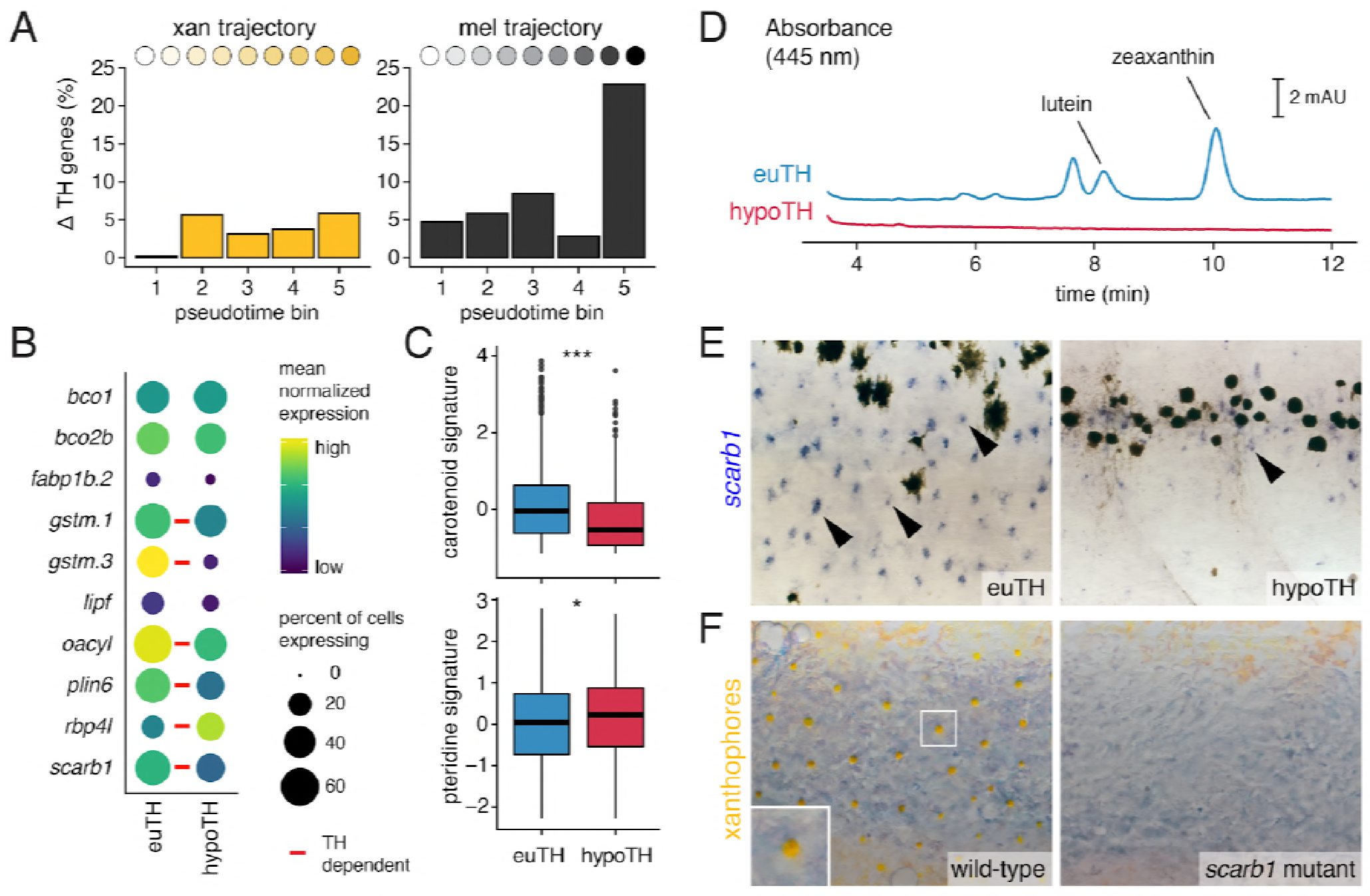
TH promotes xanthophore maturation via *scarb1*-dependent carotenoid uptake. **(A)** Proportions of differentially expressed genes in euthyroid and hypothyroid cells across pseudotime bins. Xanthophores expressed fewer TH-dependent genes than melanophores (expressed gene cutoff = 2% of bin expressing, DEGs are genes with *q*<0.05 and fold change > 1.5X). Of 160 xanthophore DEGs and 519 melanophore DEGs, only 58 were found to be overlapping. **(B)** TH-dependent expression of genes related to carotenoid pigmentation in xanthophores. Red bars: *q*<0.05, log_2_ fold-change ≥ 2.0. **(C)** Carotenoid pathway gene expression score was higher in xanthophore lineage cells of euthyroid fish compared to hypothyroid fish (*P*=1.5E-15, Wilcoxon). By contrast, pteridine pathway gene expression was marginally lower in cells from euthyroid fish (*P*=0.01). Box-and-whisker plots represent scores across groups (center line, median; box limits, upper and lower quartiles; whiskers, 1.5x interquartile range; points, outliers). **(D)** Carotenoids were detected by HPLC in skin containing xanthophores of euthyroid but not hypothyroid fish (11 SSL). **(E)** *scarb1* expression in euthyroid and hypothyroid zebrafish (10 SSL). **(F)** *scarb1* mutants lacked mature, yellow xanthophores (12 SSL).

The distinct phases of xanthophore embryonic/early larval and adult pigmentation, and the TH-dependence of the latter, led us to ask whether mechanisms underlying coloration might be stage-specific. In contrast to the defect of adult xanthophore pigmentation in *scarb1* mutants, we found that 5 dpf larval xanthophores were indistinguishable from wild-type (**Figure 6—figure supplement 4a**). Conversely, mutants lacking xanthophore pigmentation at 5 dpf have normal adult xanthophores (Odenthal et al., 1996). Because two pigment classes—carotenoids and pteridines—can contribute to xanthophore coloration, we hypothesized that visible colors at different stages depend on different pathways. Carotenoids were undetectable in euthyroid 5 dpf larvae, and carotenoid related genes were expressed at lower levels in EL xanthophores than adult xanthophores (**Figure 6—figure supplement 4B** **and C**). By contrast, pteridine pathway genes tended to be expressed similarly across stages regardless of TH status, and were even moderately upregulated in hypothyroid xanthophores (**Figure 6C**, **Figure 6—figure supplement 4C**); pteridine autofluorescence and pterinosomes were also indistinguishable between euthyroid and hypothyroid fish (**Figure 6—figure supplement 4D**; **Figure 6—figure supplement 2B**) despite the overt difference in xanthophore color with TH status [**Figure 1A**; (McMenamin et al., 2014)]. Together, these observations imply that TH induces new, carotenoid-based pigmentation, allowing transiently cryptic xanthophores to reacquire coloration during adult pattern development. TH therefore drives maturation of both xanthophores and melanophores yet has markedly different roles in each lineage, promoting re-pigmentation of the former and proliferative senescence of the latter.

### Adult pigment cell maturation programs are gated by TH receptors

Finally, to understand how TH effects are transduced in pigment cell lineages, we evaluated roles for TH nuclear receptors (TRs) that classically repress gene expression when unliganded but activate target genes when ligand (T3) is present (Brent, 2012; Buchholz et al., 2003; Hörlein et al., 1995). Genes encoding each of the three zebrafish TRs (*thraa, thrab, thrb*) were expressed by melanophores and xanthophores, yet presumptive null alleles for each unexpectedly had pigment cell complements and patterns that resembled the wild type (**Figure 7A**; **Figure 7—figure supplement 1A–D**). Given these phenotypes, we hypothesized that unliganded TRs may normally repress xanthophore differentiation. If so, we predicted that xanthophore development in hypothyroid fish should be rescued by mutation of TR. We generated fish lacking TH and TRs and found that loss of *thrab*, on its own or in conjunction with loss of *thraa*, partially restored interstripe xanthophore carotenoid deposition and reduced melanophore complements to levels indistinguishable from euthyroid fish (**Figure 7B–D; Figure 7—figure supplement 1E**). These findings suggest that unliganded repression by TRs contributes to pigment-associated phenotypes in hypothyroid fish, implying a function for TRs in repressing xanthophore repigmentation and melanophore senescence until late stages in adult pigment pattern development. Nevertheless, roles for TR are likely to be complex as we found no evidence for precocious appearance of pigment cells in euthyroid fish homozygous for *thrab* mutation, as would be expected if these receptors serve only to delay maturation to an appropriate stage (**Figure 7—figure supplement 1F**).

**Figure 7.**
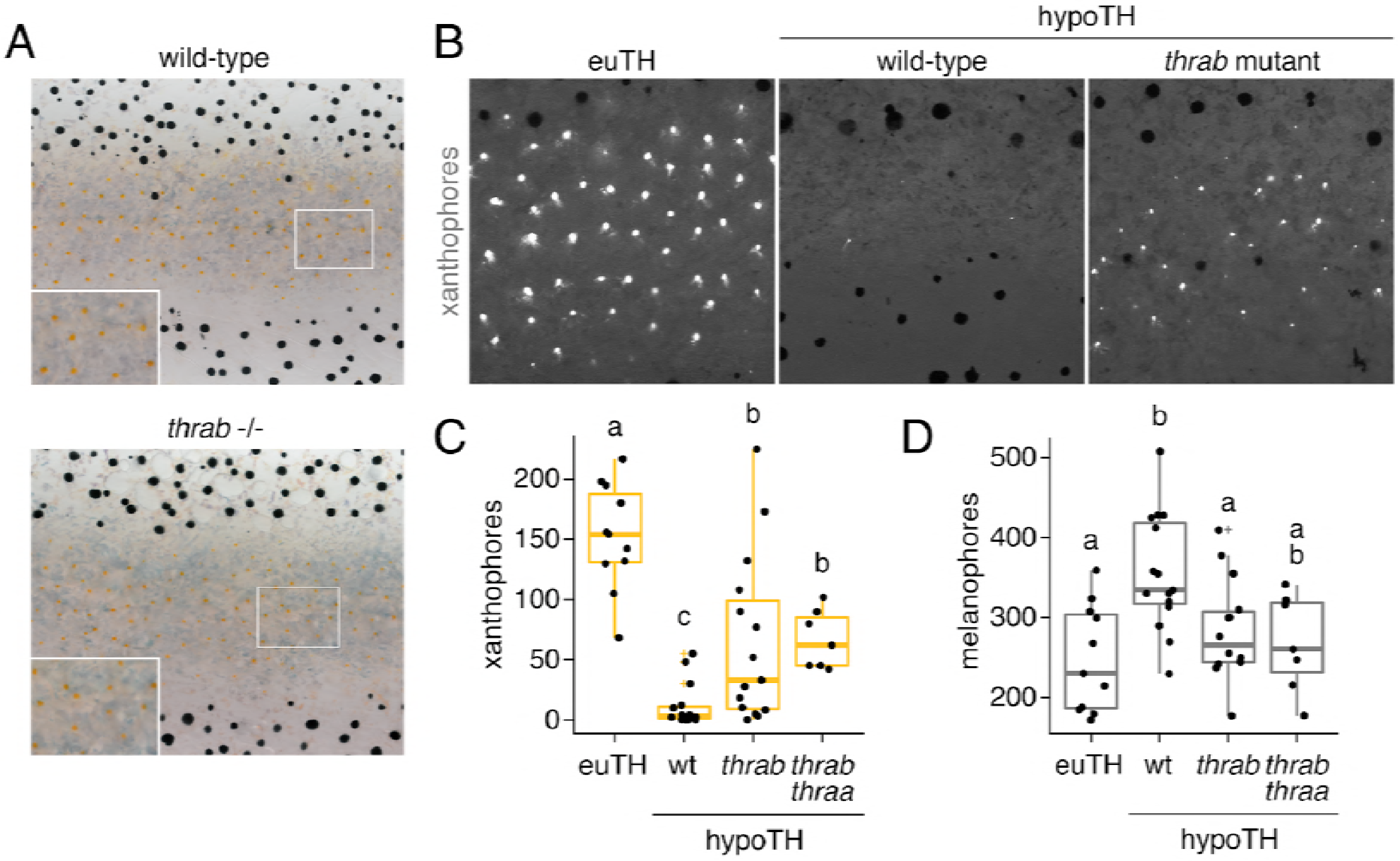
TH receptors repress developmental progression of pigment cell lineages. **(A)** TR mutants resembled wild-type; shown is *thrab*−/−. **(B)** Whereas hypothyroid fish lacked yellow-orange carotenoid-containing xanthophores, hypothyroid fish homozygous for *thrab* mutation developed substantial complements of these cells as assayed by carotenoid autofluorescence (stage-matched siblings, 11.5 SSL). **(C-D)** Homozygous *thrab* mutation partially rescued numbers of pigmented xanthophores and more fully rescued numbers of melanophores in hypothyroid fish. CRISPR/Cas9 mutagenesis of *thraa* in *thrab* −/− mutants did not significantly enhance the rescue of xanthophore maturation. Numbers of visible xanthophores and melanophores were not distinguishable between euthyroid fish wild-type or homozygous mutant for *thrab* (*P*>0.2), which are combined here. Box plots as in **Figure 6C** with different letters above data indicating significant differences in *post hoc* comparisons (Tukey HSD, *P*<0.05).

## Discussion

Our study provides insights into how TH coordinates local cellular events during the development of adult form. The stripes of adult zebrafish comprise three major classes of pigment cells that develop at specific stages and from distinct NC sublineages. Perturbations that affect the times of appearance, states of differentiation or morphogenetic behaviors of these cells can dramatically alter pattern by affecting total numbers of cells and the cascade of interactions normally required for spatial organization (Parichy and Spiewak, 2015; Patterson et al., 2014; Watanabe and Kondo, 2015). Fish lacking TH have gross defects in pigment cell numbers and pattern with ~two-fold the normal complement of melanophores and the simultaneous absence of visible xanthophores (McMenamin et al., 2014). We show that this phenotype arises not because TH normally biases cell fate specification, or has discordant effects on a particular cellular behavior that amplifies one cell type while repressing the other. Rather, our findings—combining discovery-based analyses of single-cell transcriptomic states with experiments to test specific cellular hypotheses—suggest a model whereby TH promotes maturation of both melanophores and xanthophores in distinct ways that reflect the developmental histories of these cells (**Figure 8**). Our study provides a glimpse into the diversity of cell states among post-embryonic NC-derivatives and illustrates how a single endocrine factor coordinates diverse cellular behaviors in a complex developmental process.

**Figure 8.**
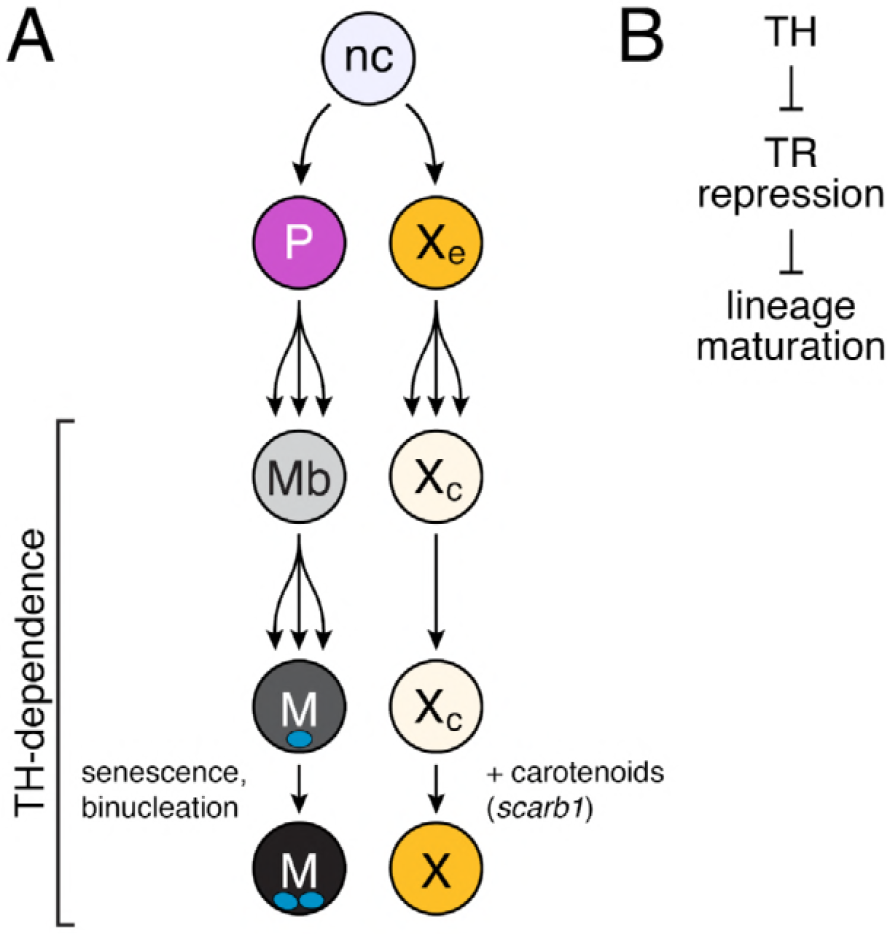
Model of TH dependence in zebrafish pigment cell lineages. **(A)** Post-embryonic progenitor-derived, specified adult melanoblasts (Mb) that expand their population and differentiate to a proliferatively arrested (McMenamin et al., 2014), senescent and binucleate state, and EL-derived cryptic xanthophores that redifferentiate as carotenoid-containing yellow/orange adult xanthophores. Disparate cell-type specific outcomes in fish lacking TH reflect differences in events required for maturation between sublineages. **(B)** TH-dependent lineage maturation involves a double negative gate, with essential repressive effects of unliganded TR.

By sampling individual cell transcriptomes across NC-derived lineages, our study complements prior investigations of lineage relationships, morphogenetic behaviors, genetic requirements, and spatial and cell-type specific gene expression profiles (Eom et al., 2015; Irion et al., 2016; Johnson et al., 1995; Kelsh et al., 2017; McMenamin et al., 2014; Parichy and Spiewak, 2015; Singh et al., 2016, 2014). Multipotent progenitors that give rise to adult melanophores, some xanthophores, and iridophores are established in the embryo and reside within peripheral nerves as development progresses (Budi et al., 2011, 2008; Dooley et al., 2013a; Singh et al., 2016; Sosa et al., 2018). As the adult pattern forms, some of these cells migrate to the hypodermis where they differentiate and integrate into dark stripes or light interstripes. The peripheral-nerve association of pigment cell progenitors in zebrafish is reminiscent of nerve-associated Schwann cell precursors that contribute to melanocytes of mammals and birds (Adameyko et al., 2009). Our collected cell-types, which include immature and mature glia, differentiating pigment cells, and presumptive progenitors of different types identify new candidate genes for promoting—and recognizing—distinct states of differentiation and morphogenetic activities, and will enable efforts to define how multipotent NC progenitors are maintained and recruited into particular lineages. That corresponding populations of embryonic and adult populations had largely overlapping transcriptomic states (**Figure 2—figure supplement 2D**) additionally highlights the intriguing problem of how specific pathways are deployed reiteratively across life cycle phases to achieve specific morphogenetic outcomes.

Our identification of a role for TH in the adult melanophore lineage illuminates how these cells develop normally and mechanisms that likely contribute to the supernumerary melanophores of hypothyroid fish. Melanoblasts derived from peripheral-nerve associated progenitors are proliferative during adult pigment pattern formation yet this activity largely ceases as the cells differentiate (Budi et al., 2011; McMenamin et al., 2014). Several lines of evidence suggest that TH promotes melanophore maturation to a senescent state: in the presence of TH, melanophores became terminally binucleate and exhibited both senescence-associated β-galactosidase activity and lysosomal content. TH also promotes the melanization of melanoblasts *ex vivo* and the proliferative arrest of melanophores *in vivo* (McMenamin et al., 2014). Our findings are broadly consistent with a role for TH in balancing proliferation and differentiation (Brent, 2012) and may be of clinical relevance, as human melanoma is associated with hypothyroidism and recurrent TH pathway mutations (Ellerhorst et al., 2003; Shah et al., 2006; Sisley et al., 1993). We suggest a model in which TH normally curtails expansion of the adult melanophore population by ensuring that cells enter a senescent state in a timely manner; in hypothyroid fish, the inappropriate retention of immature melanophores allows continued population growth during these post-embryonic stages. Effects of TH on melanophores may be direct, as these cells expressed TRs, or indirect and mediated by stromal or other cells with which melanophores interact (Lang et al., 2009). Indeed, iridophores regulate melanophore behavior during stripe establishment (Frohnhöfer et al., 2013; Patterson and Parichy, 2013) and we observed striking differences in iridophore maturation dependent on TH status (**Figure 4E**).

TH promoted the terminal differentiation of xanthophores, but in a manner distinct from melanophores. We found far fewer TH-dependent genes in xanthophores than melanophores, likely reflecting the different developmental histories of these cells. Unlike adult melanophores that arise from a transit-amplifying progenitor, most adult xanthophores develop directly from EL xanthophores that lose their pigment and then regain color late in adult pattern formation (McMenamin et al., 2014; Patterson et al., 2014) when TH levels are at a peak (Chang et al., 2012). The yellow-orange color of xanthophores can arise from pteridine pigments, carotenoid pigments, or both (Bagnara and Matsumoto, 2006; Granneman et al., 2017; Odenthal et al., 1996; Ziegler, 2003). We showed that TH positively regulates carotenoid-associated genes and carotenoid deposition, allowing cryptic xanthophores to reacquire visible pigmentation. TH did not similarly influence pteridine pathway genes. These observations suggest that TH mediates a transition from pteridine-dependent pigmentation at embryonic/early larval stages to carotenoid-dependent pigmentation of the same cells in the adult. Consistent with the notion of TH-mediated pigment-type switching, TH-dependent *scarb1* was required for carotenoid accumulation during adult pattern formation, yet mutants lacked an embryonic xanthophore phenotype. Conversely, mutants with pteridine and color deficiencies in embryonic/early larval xanthophores have normally pigmented adult xanthophores (Odenthal et al., 1996). In xanthophores, then, TH drives a state of terminal differentiation from a developmental program that is relatively more advanced than that of of progenitor-derived melanophores. That cryptic xanthophores appear poised to re-differentiate likely explains the smaller proportion of genes that were TH-dependent in these cells as compared to melanophores. Cryptic and fully differentiated xanthophores also differ from one another in their interactions with melanophores (Eom et al., 2015), and it will be interesting to determine whether genes contributing to these behaviors are under TH control as well.

Finally, our study provides clues to likely roles for TRs during adult pigment pattern formation. TR mutants lacked overt pigmentation defects yet allowed for partial rescues of both melanophore and xanthophore defects in hypothyroid fish, suggesting that unliganded TRs normally repress maturation of these lineages. Loss of TRs similarly allows the survival of congenitally hypothyroid mice (Flamant et al., 2002; Flamant and Samarut, 2003). TRs may therefore prevent the inappropriate activation of gene expression programs required for lineage maturation when TH levels are low, as is thought to occur during amphibian metamorphosis (Choi et al., 2015; Shi, 2013). Nevertheless, the failure of xanthophores to develop precociously in euthyroid fish mutant for *thrab* suggests additional requirements for TRs, or functional redundancies among their loci.

Taken together, our findings shed light on how globally available signals can control fine-grained patterning of cells within complex adult tissues.

## Materials and methods

### Staging, rearing and stocks

Staging followed (Parichy et al., 2009) and fish were maintained at ~28.5 °C under 14:10 light:dark cycles. All thyroid-ablated (Mtz-treated) and control (DMSO-treated) *Tg(tg:nVenus-v2a-nfnB)* fish were kept under TH-free conditions and were fed only *Artemia*, rotifers enriched with TH-free Algamac (Aquafauna), and bloodworms. Fish stocks used were: wild-type AB^wp^ or its derivative WT(ABb) (Eom et al., 2015); *Tg(tg:nVenus-v2a-nfnB)*^*wp.rt8*^, *Tg(aox5:palmEGFP)*^*wp.rt22*^, *Tg(tyrp1b:palm-mCherry)*^*wp.rt11*^ (McMenamin et al., 2014); *csf1ra*^*j4blue*^ (Parichy et al., 1999); *Tg(−28.5Sox10:Cre)*^*zf384*^ (Kague et al., 2012); *Tg(−3.5ubi:loxP-EGFP-loxP-mCherry)*^*cz1701*^ (Mosimann et al., 2011); *tuba8l3:nEosFP*^*vp.rt17*^, *thrab*^*vp.r31c1*^, *thraa*^*vp.r32c1*^, *thrb*^*vp.r33c1*^, *scarb1*^*vp.r32c1*^ and *tyr*^*vp.r34c1*^ (this study). Mutants and transgenic lines were maintained in the WT(ABb) genetic background. Fish were anesthetized prior to imaging with MS222 and euthanized by overdose of MS222. All procedures involving live animals followed federal, state and local guidelines for humane treatment and protocols approved by Institutional Animal Care and Use Committees of University of Virginia and University of Washington.

### Nitroreductase-mediated cell ablation

To ablate thyroid follicles of *Tg(tg:nVenus-2a-nfnB)*, we incubated 4 day post-fertilization (dpf) larvae for 8 h in 10 mM Mtz with 1% DMSO in E3 media, with control larvae incubated in 1% DMSO in E3 media. For all thyroid ablations, treated individuals were assessed for loss of nuclear-localizing Venus (nVenus) the following day. Ablated thyroid glands fail to regenerate (McMenamin et al., 2014) and absence of regeneration in this study was confirmed by continued absence of nVenus expression.

### Mutant and transgenic line production

For CRISPR/Cas9 mutagenesis, 1-cell stage embryos were injected with 200 ng/μl sgRNAs and 500 ng/μl Cas9 protein (PNA Bio) using standard procedures (Shah et al., 2015). Guides were tested for mutagenicity by Sanger sequencing and injected fish were reared through adult stages at which time they were crossed to *Tg(tg:nVenus-v2a-nfnB)* to generate heterozygous F1s from which single allele strains were recovered. CRISPR gRNA targets (excluding protospacer adjacent motif) are included in **Table 7**. Mutant alleles of *scarb1* and TR loci are provided in **Figure 6—figure supplement 3** and **Figure 7—figure supplement 1**, respectively. The melanin free *tyr*^*vp.r34c1*^ allele generated for analyses of melanophore lysosomal content exhibits a 4 nucleotide deletion beginning at position 212 that leads to novel amino acids and a premature stop codon (H71QEWTIESDGL*).

To label nuclei of adult melanophores, BAC CH73-199E17 containing the *puma* gene *tuba8l3 (Larson et al., 2010)* was recombineered to contain nuclear-localizing photoconvertible fluorophore EosFP using standard methods (Sharan et al., 2009; Suster et al., 2011).

### Imaging

Images were acquired on: Zeiss AxioObserver inverted microscopes equipped with Axiocam HR or Axiocam 506 color cameras; a Zeiss AxioObserver inverted microscope equipped with CSU-X1 laser spinning disk (Yokogawa) and Orca Flash 4.0 camera (Hamamatsu Photonics); or a Zeiss LSM 880 scanning laser confocal microscope with Fast Airyscan and GaAsP detectors. Images were corrected for color balance and adjusted for display levels as necessary with conditions within analyses treated identically.

### Cell counts

Melanophores and xanthophores were counted within regions defined dorsally and ventrally by the margins of the primary stripes, anteriorly by the anterior margin of the dorsal fin, and posteriorly by five myotomes from the start. Only hypodermal melanophores were included in analysis; dorsal melanophores and those in scales were excluded. Mature xanthophores were counted by the presence of autofluorescent carotenoid with associated yellow pigment. Cell counts were made using ImageJ. Individual genotypes of fish assessed were confirmed using PCR or Sanger sequencing.

### *In situ* hybridization

*In situ* hybridization (ISH) probes and tissue were prepared as described (Quigley et al., 2004). Probes were hybridized for 24 hr at 66°C. Post-hybridization washes were performed using a BioLane HTI 16Vx (Intavis Bioanalytical Instruments), with the following parameters: 2x SSCT 3 × 5 min, 11 × 10 min at 66°C; 0.2x SSCT 10 × 10 min; blocking solution [5% normal goat serum (Invitrogen), 2 mg/mL BSA (RPI) in PBST] for 24 hr at 4 °C; anti-Dig-AP, Fab fragments (1:5000 in blocking solution, Millipore-Sigma) for 24 hr at 4 °C; PBST 59 × 20 min. AP staining was performed as described (Quigley et al., 2004).

### Pigment analyses

Xanthophore pigments were examined by imaging autofluorescence in eGFP and DAPI spectral ranges for carotenoids and pteridines respectively. For imaging pteridines, fish were euthanized and treated with dilute ammonia to induce autofluorescence (Odenthal et al., 1996).

For analyses of carotenoid contents by HPLC we pooled three skin samples from each genotype and condition (Mtz-treated or control) into two separate samples. We homogenized the tissue in a glass dounce homogenizer with 1 ml of 0.9% sodium chloride and quantified the protein content of each sample with a bicinchoninic acid (BCA) assay (23250, Thermo). We then extracted carotenoids by combining the homogenates with 1 ml methanol, 2 ml distilled water, and 2 ml of hexane:*tert*-methyl butyl ether (1:1 vol:vol), separated the fractions by centrifuging, collected the upper solvent fraction, and dried it under a stream of nitrogen. We saponified these extracts with 0.2 M NaOH in methanol at room temperature for four hours following the protocol described in (Toomey and McGraw, 2007). We extracted the saponified carotenoids from this solution with 2 ml of hexane:*tert*-methyl butyl ether (1:1 vol:vol) and dried the solvent fraction under a stream of nitrogen. We resuspended the saponified extracts in 120 μl of methanol:acetonitrile 1:1 (vol:vol) and injected 100 µl of this suspension into an Agilent 1100 series HPLC fitted with a YMC carotenoid 5.0 µm column (4.6 mm × 250 mm, YMC). We separated the pigments with a gradient mobile phase of acetonitrile:methanol:dichloromethane (44:44:12) (vol:vol:vol) through 11 minutes, a ramp up to acetonitrile:methanol:dichloromethane (35:35:30) for 11-21 minutes and isocratic conditions through 35 minutes. The column was warmed to 30°C, and mobile phase was pumped at a rate of 1.2 ml min^−1^ throughout the run. We monitored the samples with a photodiode array detector at 400, 445, and 480 nm, and carotenoids were identified and quantified by comparison to authentic standards (a gift of DSM Nutritional Products, Heerlen, The Netherlands). Analyses of 5 dpf wild-type and *csf1ra* mutants used only larval heads where xanthophores are abundant in the wild type; other procedures were the same as for later stages.

### Immunohistochemistry and Oil-red-O staining

Skins of *Tg(aox5:palmEGFP)* euthyroid and hypothyroid zebrafish (8.6–10.4 SSL) were dissociated and plated at low density in L-15 medium (serum free) on collagen-coated, glass bottom dishes (Mattek) for 5 h. Cells were then fixed with freshly prepared 4% PFA for 15 m, rinsed with PBST (0.1%), blocked (5% goat serum, 1% BSA, 1X PBS), then incubated at 4°C overnight with rabbit anti-GFP primary antibody (ThermoFisher). Stained cells were rinsed 3X with 1X PBS and fixed again with 4% PFA for 30 minutes. Cells were then rinsed twice with ddH2O, washed with 60% isopropanol for 5 min, and then dried completely. Cells were incubated with filtered, Oil Red O solution (5 mM in 60% isopropanol) for 10 min, and rinsed 4X with ddH20 before imaging (Koopman et al., 2001). All GFP+ cells were imaged across two plates per condition and were scored for presence or absence of red staining.

### Cellular senescence assays

For assaying senescence of melanophores *ex vivo*, skins from euthyroid and hypothyroid fish (*n*=3 each, 11 SSL) were dissociated and plated on glass-bottom, collagen coated dishes (MatTek) in L-15 medium (Gibco) and incubated overnight at 28°C. Cells were then rinsed with dPBS, fixed and stained using a Senescence β-;Galactosidase Staining Kit (Cell Signaling Technologies, cat. #9860) according to manufacturer’s instructions (Ceol et al., 2011; Dimri et al., 1995). Staining was carried out for 48 h at pH 6 prior to imaging.

To assay senescence as measured by lysosomal content (Kurz et al., 2000; Lee et al., 2006) of melanophores by FACS skins from euthyroid and hypothyroid *Tg(tyrp1b:palm-mCherry; tuba8l3:nEOS*), *tyr* fish lacking melanin (*n*=12 each) were dissociated and resuspended 1% BSA / 5% FBS / dPBS. Cells were incubated for 1 h with Lysotracker (75 nM) (ThermoFisher, L12492) and Vybrant DyeCycle Violet stain (5 μM) (ThermoFisher, V35003) shaking at 500 rpm, 28°C. Without washing, cells were FAC sorted. Single transgene controls and wild type cells were used to adjust voltage and gating. Prior to analysis of fluorescence levels, single cells were isolated by sequentially gating cells according to their SSC-A vs. FSC-A, FSC-H vs FSC-W and SSC-H vs SSC-W profiles according to standard flow cytometry practices. Intact live cells were then isolated by excluding cells with low levels of DyeCycle violet staining (DAPI-A). As expected these cells express a wide range of our *tuba8l3:nlsEosFP* transgene as determined by levels of green fluorescence (FITC-A). Melanophores were isolated by identifying cells with high fluorescence in the FITC-A and mCherry-A channels which describe expression of the *tuba8l3:nlsEosFP* and *tyrp1b:palm-mCherry* transgenes. Lastly, lysosomal content of melanophores was determined by the median fluorescence intensity of our lysosomal marker, Lysotracker Deep Red (APC-A) (**Figure 4—figure supplement 1**). The data were collected on a FACS ARIA using FACSDiva version 8 software (BD Biosciences).

Data were analyzed using FlowJo v10.

### Transmission electron microscopy

Fish were euthanized then fixed in sodium cacodylate buffered 4% glutaraldehyde overnight at 4°C. Trunk regions were dissected then tissue stained in 2% osmium tetroxide for 30 minutes, washed, and then stained in 1% uranyl acetate overnight at 4°C. Samples were dehydrated with a graded ethanol series then infiltrated with a 1:1 propylene oxide:Durcupan resin for 2 hours followed by fresh Durcupan resin overnight and flat embedded prior to polymerization. Blocks were thin sectioned on a Leica EM UC7 and sections imaged on a JEOL 1230 transmission electron microscope.

### Tissue dissociations and FACS

Trunks or skins of staged, post-embryonic zebrafish (7.2–11.0 SSL) were dissected (*n*=8 per replicate) and enzymatically dissociated with Liberase (Sigma-Aldrich cat. 5401119001, 0.25 mg/mL in dPBS) at 25°C for 15 min followed by manual trituration with a flame polished glass pipette for 5 min. Cell suspensions were then filtered through a 70 μm Nylon cell strainer to obtain a single cell suspension. Liberated cells were re-suspended in 1% BSA / 5% FBS in dPBS and DAPI (0.1 μg/mL, 15 min) before FACS purification. All plastic and glass surfaces of cell contact were coated with 1% BSA in dPBS before to use. Prior to sorting for fluorescence levels, single cells were isolated by sequentially gating cells according to their SSC-A vs. FSC-A, FSC-H vs FSC-W and SSC-H vs SSC-W profiles according to standard flow cytometry practices. Cells with high levels of DAPI staining were excluded as dead or damaged. Cells from wild-type and *Tg(ubi:switch*) zebrafish without Cre were used as negative control to determine gates for detection of mCherry and GFP fluorescence, then cells from *Tg(sox10:Cre; ubi:switch)* zebrafish were purified according to these gates. NC-derived cells cells were isolated by identifying cells with high fluorescence in the mCherry-A channel which describes expression of the *ubi:loxP-EGFP-loxP-mCherry* transgene after permanent conversion to *ubi:mCherry* after exposure to *Sox10:Cre* (see **Figure 1—figure supplement 1C**). All samples were kept on ice except during Liberase incubation, and sorted chilled.

### RT-PCR

Skin tissue from stage-matched fish was dissociated as above and melanophores and xanthophores were FAC sorted for the presence *aox5:palmeGFP* or *tyrp1b:palm-mCherry*, respectively. RNA was extracted from pools of 1000 cells using the RNAqueous-Micro kit (Thermo Fisher, cat. AM1912). Full length cDNA was synthesized with Superscript III reverse transcriptase (Thermo Fisher, cat. #18080093). Amplifications were 40 cycles with Q5 DNA polymerase (NEB, M0492), 38 cycles at 94°C, 30s; 67°C, 20s; 72°C, 20s. For primer sequences (*actb1, thraa, thrab, thrb*), see **Table 7.**

### Single cell collection, library construction and sequencing

Whole-trunks or skins were collected from stage-matched *Tg(tg:nVenus-2a-nfnB)* euthyroid and hypothyroid siblings, dissociated, and *sox10*:Cre:mCherry+ cells isolated by FACS.

We replicated the experiment three times. For each replicate, we collected cells from euthyroid and hypothyroid fish at 7.2 SSL, 8.6 SSL, and 9.6 SSL (mid-larval, 6–10 fish per stage, per replicate) and sorted equal numbers of mCherry+ cells from each group into a single sample. Cells were pelleted and resuspended in 0.04% ultrapure BSA (ThermoFisher Scientific). Representing a terminal stage of pigment pattern development, we also collected mCherry+ cells from one sample within each replicate of 11 SSL (juvenile, 5 fish per condition) euthyroid and hypothyroid fish. To capture cells representing the EL pigment pattern, we collected mCherry+ cells from 5 dpf larvae (50 fish). In each experiment, we ran parallel euthyroid and hypothyroid samples (fish were siblings). For each sample, we targeted 2000–4000 cells for capture using the Chromium platform (10X Genomics) with one lane per sample. Single-cell mRNA libraries were prepared using the single-cell 3’ solution V2 kit (10X Genomics). Quality control and quantification assays were performed using a Qubit fluorometer (Thermo Fisher) and a D1000 Screentape Assay (Agilent). Libraries were sequenced on an Illumina NextSeq 500 using 75-cycle, high output kits (read 1: 26 cycles, i7 Index: 8 cycles, read 2: 57 cycles). Each sample was sequenced to an average depth of 150 million total reads. This resulted in an average read depth of ~40,000 reads/cell after read-depth normalization.

### scRNA-Seq data processing

We found that for many genes, annotated 3’ UTRs in the Ensembl 93 zebrafish reference transcriptome were shorter than true UTR lengths observed empirically in pileups of reads mapped to the genome. This led to genic reads being counted as intergenic. To correct for this bias in aligning reads to the transcriptome, we extended all 3’ UTR annotations by 500 bp. In rare cases, UTR extension resulted in overlap with a neighboring gene and in these instances we manually truncated the extension to avoid such overlap. We built a custom zebrafish STAR genome index using gene annotations from Ensembl GRCz11 with extended 3’ UTRs plus manually annotated entries for mCherry transcript, filtered for protein-coding genes (with Cell Ranger *mkgtf* and *mkref* options). Final cellular barcodes and UMIs were determined using Cell Ranger 2.0.2 (10X Genomics) and cells were filtered to include only high-quality cells. Cell Ranger defaults for selecting cell-associated barcodes versus barcodes associated with empty partitions were used. All samples were aggregated (using 10X Cell Ranger pipeline “cellranger aggr” option), with intermediary depth normalization to generate a gene-barcode matrix containing ~25,000 barcoded cells and gene expression counts.

### UMAP visualization and clustering

We used Uniform Manifold Approximation and Projection (UMAP) (McInnes et al., 2018) to project cells in two or three dimensions and performed louvain clustering (Blondel et al., 2008) using the reduceDimension and clusterCells functions in Monocle (v.2.99.1) using default parameters (except for, reduceDimension: reduction_method=UMAP, metric=cosine, n_neighbors=30, mid_dist=0.5; clusterCells: res=1e-3, k=15). We assigned clusters to cell types based on the detection of published marker genes. Cells isolated from euthyroid and hypothyroid fish were combined to maintain consistency of analysis and for comparisons between groups. Batch correction methods were not used between the two groups or across samples because we did not observe sample-specific separation or clustering in UMAP space. Cells with more than 15,000 UMIs were discarded as possible doublets. All genes were given as input to Principal Components Analysis (PCA). The top 30 principal components (high-loading, based on the associated scree plot) were then used as input to UMAP for generating either 2D or 3D projections of the data. For, subclustering of pigment cell clusters (melanophores, iridophores, xanthophores, and pigment progenitors) we subsetted the data set and again applied UMAP dimensionality reduction and louvain clustering.

### Differential expression analysis to determine cell-type markers

To identify genes expressed cell-type specifically, we used the principalGraphTest function in Monocle3 (v.2.99.1) with default parameters (Cao et al., 2018). This function uses a spatial correlation analysis, the Moran’s I test, to assess spatially restricted gene expression patterns in low dimensional space. We selected markers by optimizing for high specificity, expression levels and effect sizes within clusters (For extended list of cell-type specific genes, see **Table 1**).

### Trajectory analysis

The top 800 highly dispersed genes (**Table 5**) within euthyroid pigment cells (melanophores, xanthophores, iridophores, and pigment progenitors) were chosen as feature genes to resolve pseudotemporal trajectories using the setOrderingFilter, reduceDimension, and orderCells functions in Monocle (v2.9.0) using default parameters with the exception of setting max_components = 3 and num_dim = 10 to generate the trajectory in 3D with the top 10 PCs (high-loading based on scree plot) during dimensionality reduction.

### Branched Expression Analysis Modeling (BEAM)

After running trajectory analysis on pigment cells, we used the BEAM function in Monocle (v.2.9.0) with default settings (except, branch_point = 3) to determine differentially expressed genes between trajectory branches. To generate the BEAM heatmap for the three pigment cell trajectory branches, we used the plot_multiple_branches_heatmap function with default settings (except assigning branch 1, 5, and 6 to iridophores, melanophores, and xanthophores, respectively; and num_clusters = 6). Genes were selected by significance levels for the three-branch BEAM analysis with additional significant genes added from the melanophore and iridophore two-branch analysis for more even distribution of genes across lineages (*q*<6.0E-11 for all genes, except for *pax3a* (starred, *q*=0.03) which is a positive indicator of early pseudotime for all lineages).

### Differential expression analysis across pseudotime

To determine differentially expressed genes over pseudotime that were TH-dependent, we filtered the data set for genes expressed in at least 5 cells and performed differential expression analysis using a full model of sm.ns(Pseudotime, df=3)*condition and a reduced model of sm.ns(Pseudotime, df=3).

### Development and analysis of pathway signature scores

Gene sets for signature scores were selected using gene ontology (terms and gene sets from zfin.org; cell-cycle, unfolded protein response, AP-1 transcription factor complex members) or manual curation based on literature when required (carotenoid, pteridine, melanin) (see **Table 4**). Signature scores were calculated by generating z-scores (using scale( )) of the the mean of expression values (log transformed, size factor normalized) from genes in a given set.

### Statistics

Parametric, non-parametric and multiple logistic regression analyses were performed using JMP 14.0 (SAS Institute, Cary, NC) or *R* [version 3.5.0] (R Core Team, 2017). For parametric analyses, residuals were assessed for normality and homoscedasticity to meet model assumptions and no transformations were found to be warranted.

### Data availability

Data is available on GEO via accession GSEXXXXXX. (Data will be made publically available upon publication).

### Code availability

Code will be made available to reviewers upon request.

## Acknowledgments

For assistance we thank D. Jackson, D. Huang, D.S. Eom, D. Raible, A. Fulbright, A. Leith, A. Schwindling, T. Linbo, D. White, and E. Parker.

---

**Figure 2—figure supplement 1.**
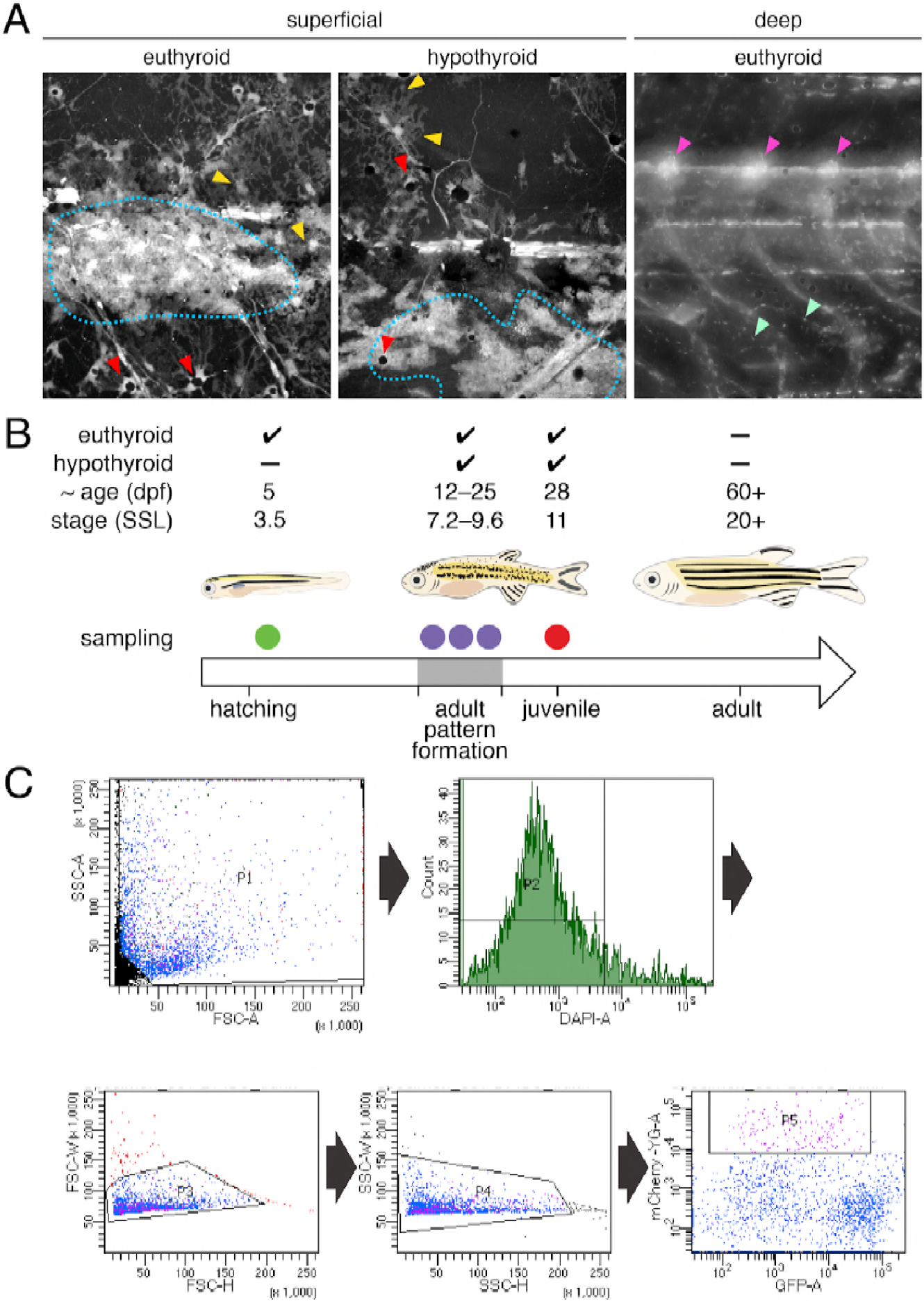
Experimental design and isolation of NC-derived cells from post-embryonic zebrafish. **(A)** Fish transgenic for *sox10:cre* and *ubi:loxP-EGFP-loxP-mCherry* permanently and robustly expressed mCherry in NC-derived cells of both euthyroid and hypothyroid fish(Kague et al., 2012; Mosimann et al., 2011). At superficial layers, mCherry+ xanthophores (yellow arrowheads), melanophores (red arrowheads), and iridophores (blue dotted line) were apparent. At deeper layers, mCherry+ cells were found in dorsal root ganglia (magenta arrowheads) and other locations (e.g., mint arrowheads), potentially representing glia, neurons, progenitors and other cell types. mCherry+ cells of non-NC origin were evident as well (see Extended data Fig. 2). Stage shown is 9.8 (mm) standardized standard length (SSL) (Parichy et al., 2009). **(B)** Single-cell RNA-Seq (scRNA-Seq) experimental design. To ensure that progenitors, cells at intermediate states of specification and commitment, and fully differentiated cells were captured, euthyroid and hypothyroid fish were collected at a range of stages encompassing adult pattern formation (7.2–9.8 SSL) and from juvenile fish (11 SSL) in which the first two adult stripes had fully formed (Parichy et al., 2009). To compare transcriptomic signatures of NC-derived cells from embryonic–early larval and middle larval–juvenile stages, cells were additionally collected from euthyroid larvae at 5 dpf (3.5 SSL). **(C)** Cells were FACS-isolated by detection of mCherry vs. GFP fluorescence. **(D)** Representative FAC sort for NC-derived cells from post-embryonic skins and trunks. Single cells were isolated by sequentially gating cells according to their SSC-A vs. FSC-A, FSC-H vs. FSC-W, and SSC-H vs. SSC-W profiles according to standard flow cytometry practices. Cells with high levels of DAPI staining were excluded as dead or damaged. NC-derived cells were isolated by identifying cells with high fluorescence in the mCherry-A channel which describes expression of the ubi:loxP-EGFP-loxP-mCherry transgene after permanent conversion to ubi:mCherry after exposure to *sox10*:Cre (see **Figure 2—figure supplement 1A**).

**Figure 2—figure supplement 2.**
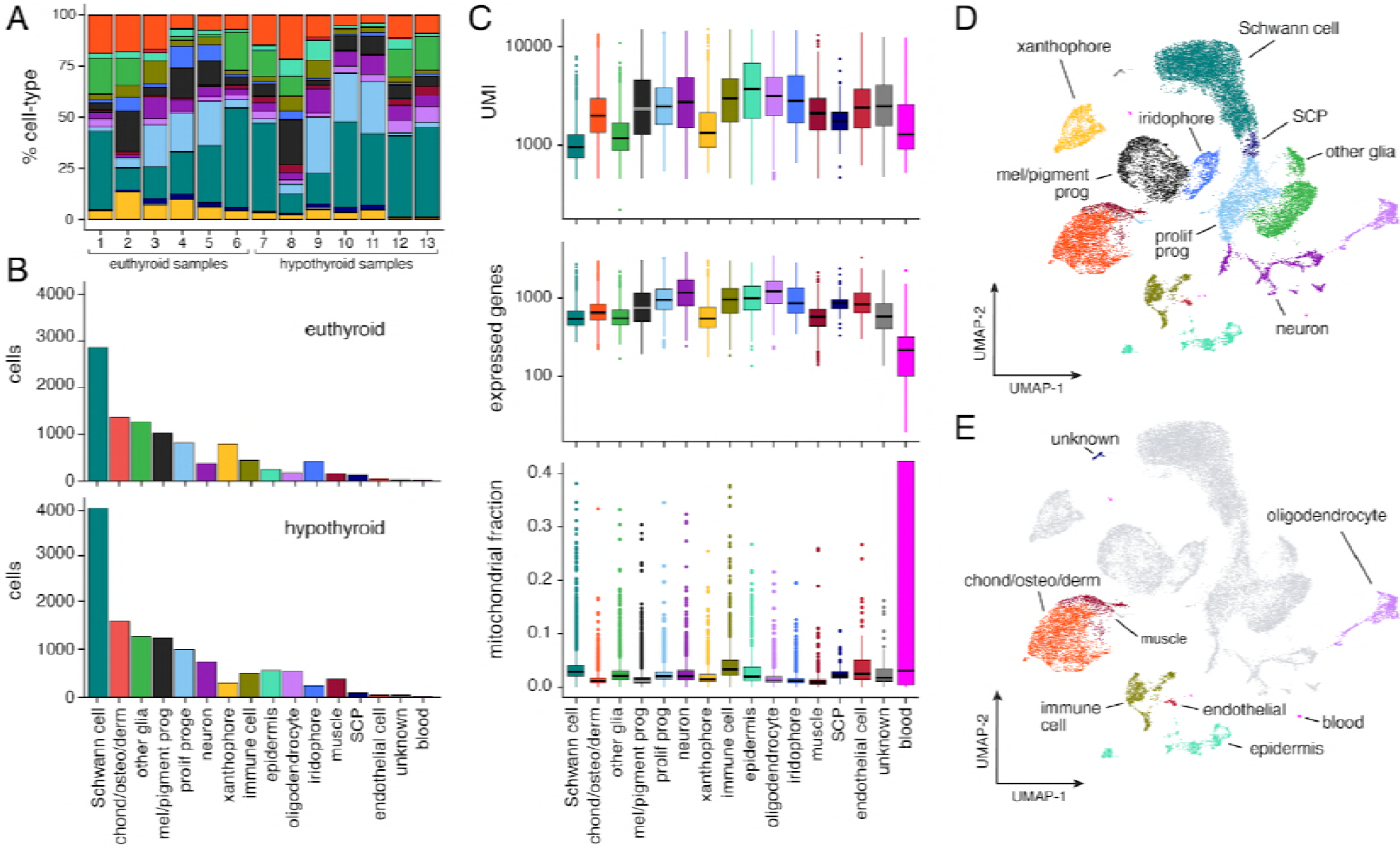
Population characteristics for full scRNA-Seq dataset from post-embryonic zebrafish. **(A)** Percentages of cell types across samples (for details, see **Table 3**). All cell types are represented in each sample in similar overall proportions. Colors correspond to cell types in (B–E). **(B)** Counts of cells by type captured from euthyroid and hypothyroid fish at post-embryonic stages (≥7.2 SSL). **(C)** Counts of unique molecular identifiers (UMI) and unique genes expressed, as well as fractions of mitochondrial reads by cell-type (shown are medians with boxes spanning interquartile ranges; vertical lines indicate farthest observations of data with outlier samples shown individually). Increased fractions of mitochondrially encoded genes may indicate broken cell (Ilicic et al., 2016); consistent with this idea, Schwann cells—many of which are expected to be damaged owing to their concentrically layered morphology—exhibited one of the largest overall proportions of UMIs derived from mitochondrial genes. **(D)** Two-dimensional UMAP representation of all post-embryonic cells (≥7.2 SSL) captured in scRNA-Seq that passed quality thresholds (22,613 cells, 13 combined samples). **(E)** Same cells as in D, with presumptive NC-derivatives shown in grey and presumptive non-NC derived cells highlighted by type.

**Figure 2—figure supplement 3.**
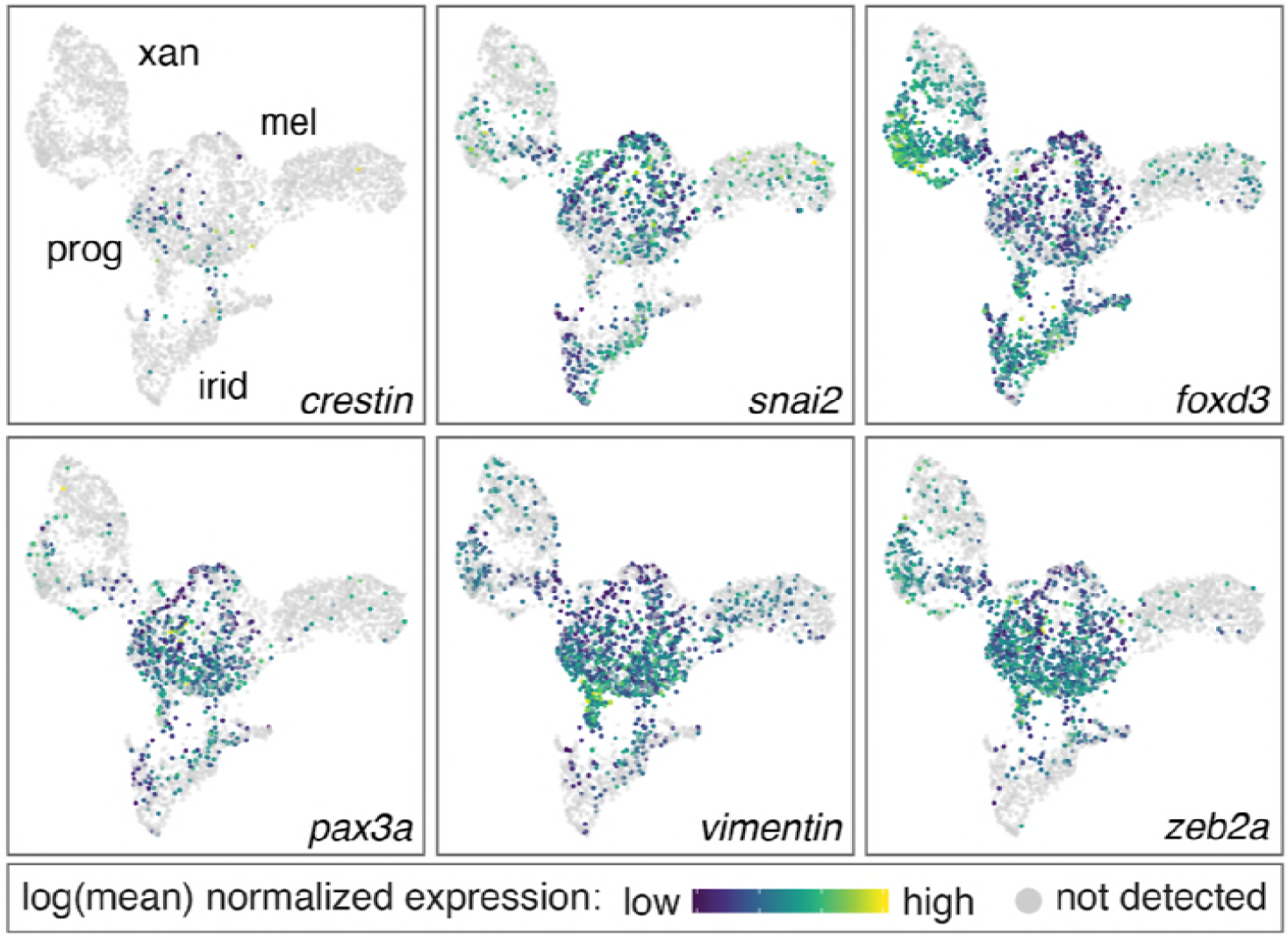
Genes enriched in pigment progenitor clusters include known markers of embryonic NC cells. Genes enriched in the pigment progenitor cluster compared to differentiated pigment cells included loci expressed in embryonic, migratory NC cells of zebrafish and other species (*crestin, snai2, foxd3, pax3a, vim, zeb2a*) (Kaufman et al., 2016; Kelsh et al., 2000a; Luo et al., 2001; Minchin and Hughes, 2008; Thisse et al., 1995; Van de Putte et al., 2003; Ziller et al., 1983). Color indicates relative expression of each gene within individual cells (xan, xanthophores; mel, melanophores; irid, iridophores; prog, pigment progenitors).

**Figure 2—figure supplement 4.**
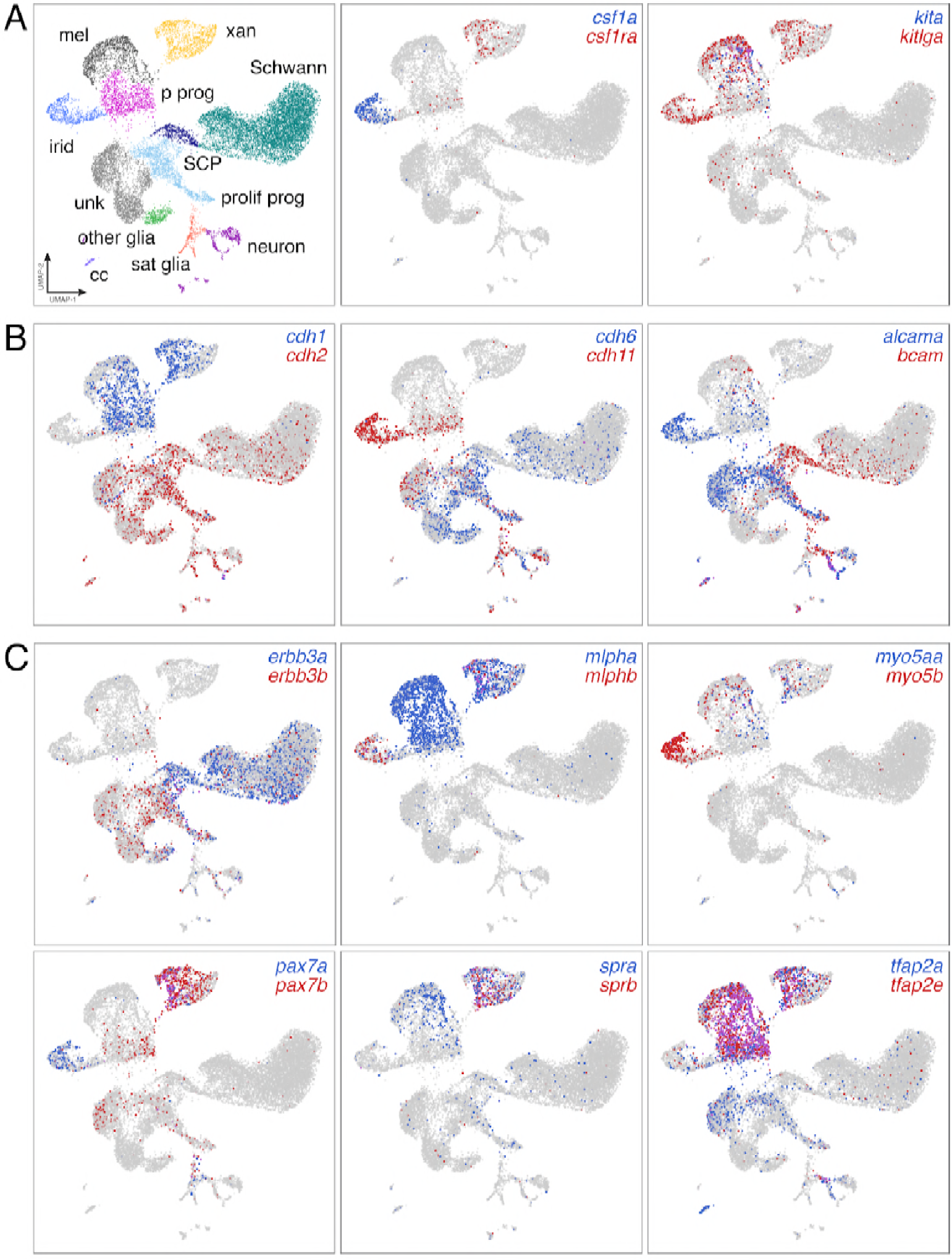
Distinct domains of gene expression across diverse NC derivatives. **(A)** UMAP representation of major NC derivatives (left) with coordinate expression of two exemplar ligand–receptor gene pairs (middle and right). Xanthophores are known to express and require type III receptor tyrosine kinase (RTK) gene *colony stimulating factor 1 receptor a* (*csf1ra*) and to depend for their development on *csf1a* expressed by nearby iridophores (Parichy et al., 2000b; Patterson and Parichy, 2013). Melanophores require the type III RTK gene *kita (Parichy et al., 1999)* and ligand encoded by *kitlga*, expressed by skin(Hultman et al., 2007; Patterson and Parichy, 2013), but also pigment cells and progenitors revealed here by scRNA-Seq. **(B)** Pairs of cell adhesion molecules with distinct expression domains suggest differing morphogenetic requirements between pigment cell and other lineages (*cdh1* vs. *cdh2*), and for specific cell types [e.g., *cdh11* of iridophores, which form epithelium-like mats within adult interstripes (Singh et al., 2014)]. **(C)** In teleosts, an ancient clade-specific genome duplication resulted in extra genes, allowing for subfunctionalization and retention of some paralogs (Braasch et al., 2015, 2009). scRNA-Seq revealed different degrees to which paralog expression has been partitioned across NC derived cell types. For example, proliferating progenitors and unknown (unk) cells were more likely to express receptor tyrosine kinase gene *erbb3b*, required for development of glia and adult melanophores(Budi et al., 2008), whereas Schwann cells and Schwann cell progenitors (SCP) were more likely to express *erbb3a*. Likewise, xanthophores express and require transcription factor genes *pax7a* and *pax7b (Nord et al., 2016)*, but iridophores were also marked by *pax7a* expression, suggesting the possibility of functional significance to this cell type exclusive of *pax7b* activities (A–C, expression thresholds=1.)

**Figure 2—figure supplement 5.**
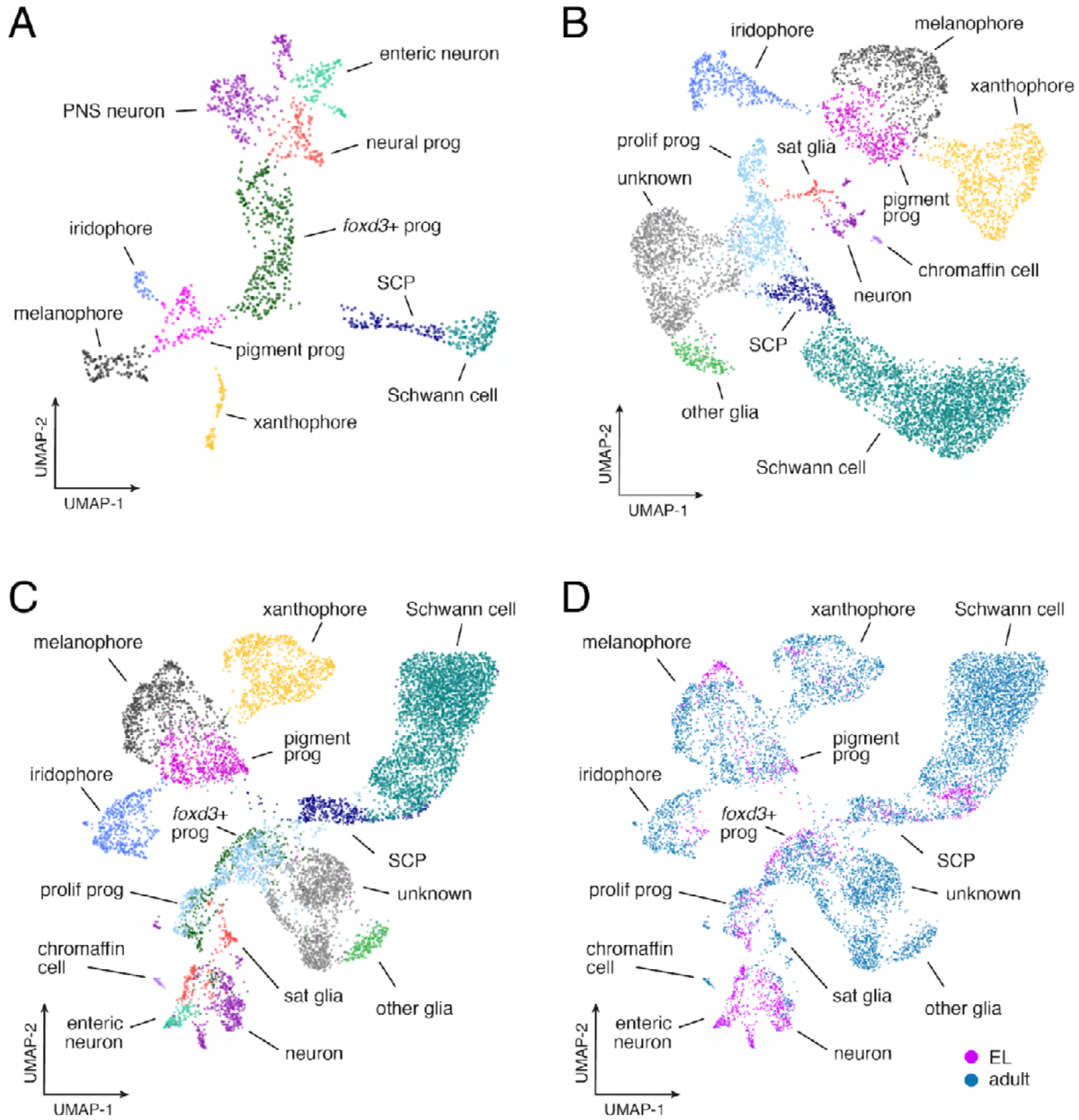
Similarities and differences between EL and adult gene expression programs. **(A)** UMAP representation of EL NC-derived cells (*n*=1,466) isolated from euthyroid 5 dpf fish (3.5 SSL; prog, progenitor; SCP, Schwann cell precursors; PNS, non-enteric peripheral nervous system). **(B)** Adult NC-derived cells (*n*=7,611) isolated from euthyroid middle larva–juvenile (≥7.2 SSL) fish (sat, satellite). **(C)** Combined EL and adult. **(D)** Comparison of EL (magenta) and adult (blue-green) cell distributions revealed broadly overlapping domains of xanthophores in UMAP space, consistent with known derivation of most adult xanthophores from EL xanthophores (McMenamin et al., 2014) [and see Main text].

**Figure 3—figure supplement 1.**
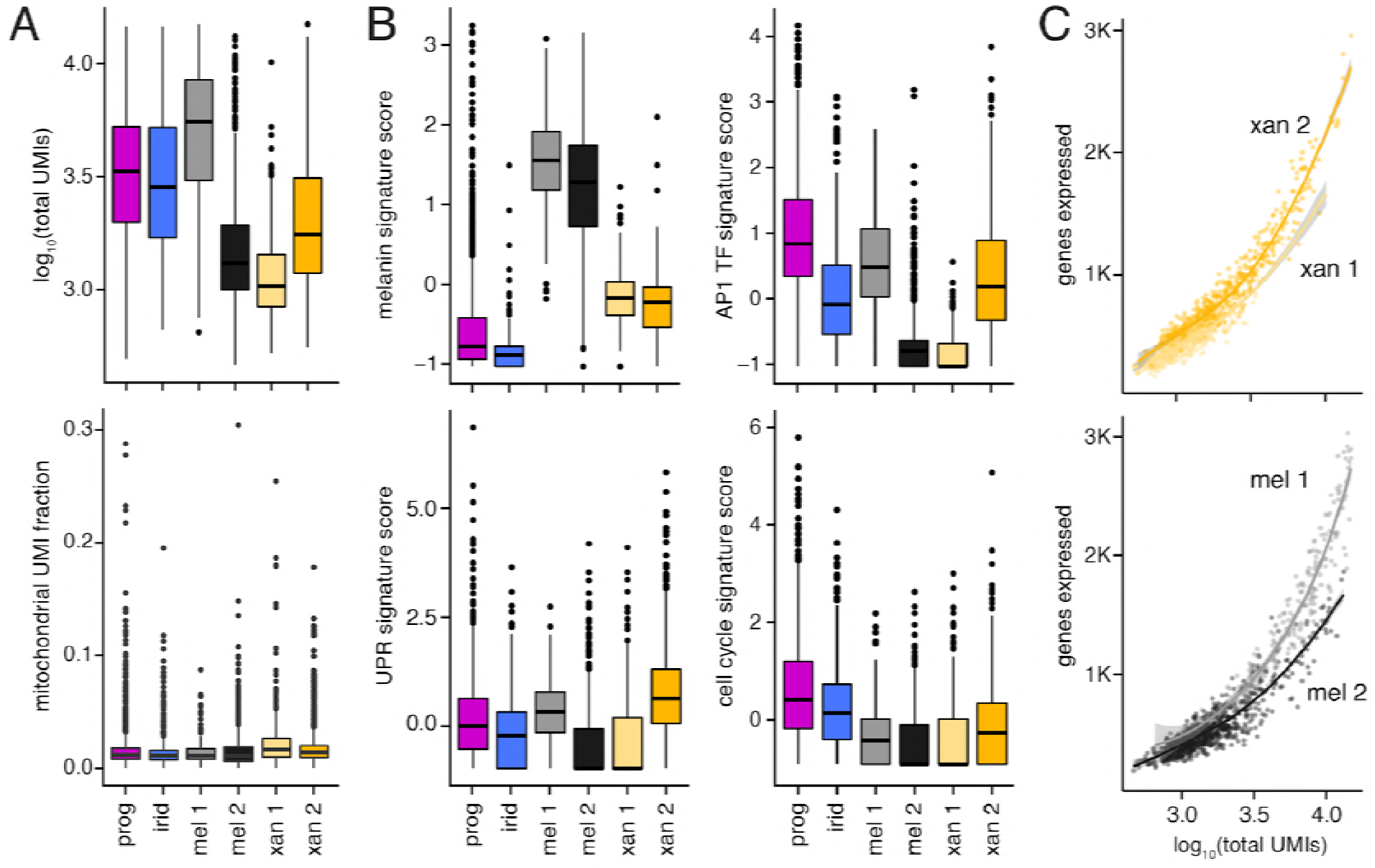
Differences between melanophore and xanthophore sub-populations revealed distinct levels and types of transcriptional activity. **(A)** Median transcript numbers (unique molecular identifiers, UMIs; upper plot) differed across pigment cell subpopulations. Reduced total RNA content is associated with a G0 cell state (i.e., quiescence, replicative senescence, and/or terminal differentiation) (Coller et al., 2006; Darzynkiewicz et al., 1980); low median UMI counts in sub-clusters of melanophores and xanthophores (mel 2, xan 1) suggest alternative states within these differentiated cell populations. Mitochondrial read fraction (lower plot) was low and consistent across sub-clusters, suggesting that low median UMI counts (upper plot) were unlikely to reflect damage specifically incurred by particular cell types(Ilicic et al., 2016). **(B)** Both melanophore subclusters highly express genes associated with melanin synthesis, indicating that they are both represent true melanophores. However, cell clusters with lower median UMIs (mel 2, xan 1) exhibited gene expression trends indicative of curtailed transcriptional and translational activity, including reduced expression of AP-1 transcriptional complex members (AP1 TF signature score), and genes involved in unfolded protein response (UPR signature score), and proliferation (cell-cycle signature score) (Chinenov and Kerppola, 2001; Patil and Walter, 2001; Riabowol et al., 1992) (for details of genes in signature scores, see **Table 4**). **(C)** Melanophores and xanthophores in subclusters with lower total UMI counts expressed fewer unique genes compared to cells in the other subcluster regardless of equivalent UMI counts, consistent with a more restricted gene expression profile of cells in a terminally differentiated state (shaded areas indicate standard error bounds).

**Figure 3—figure supplement 2.**
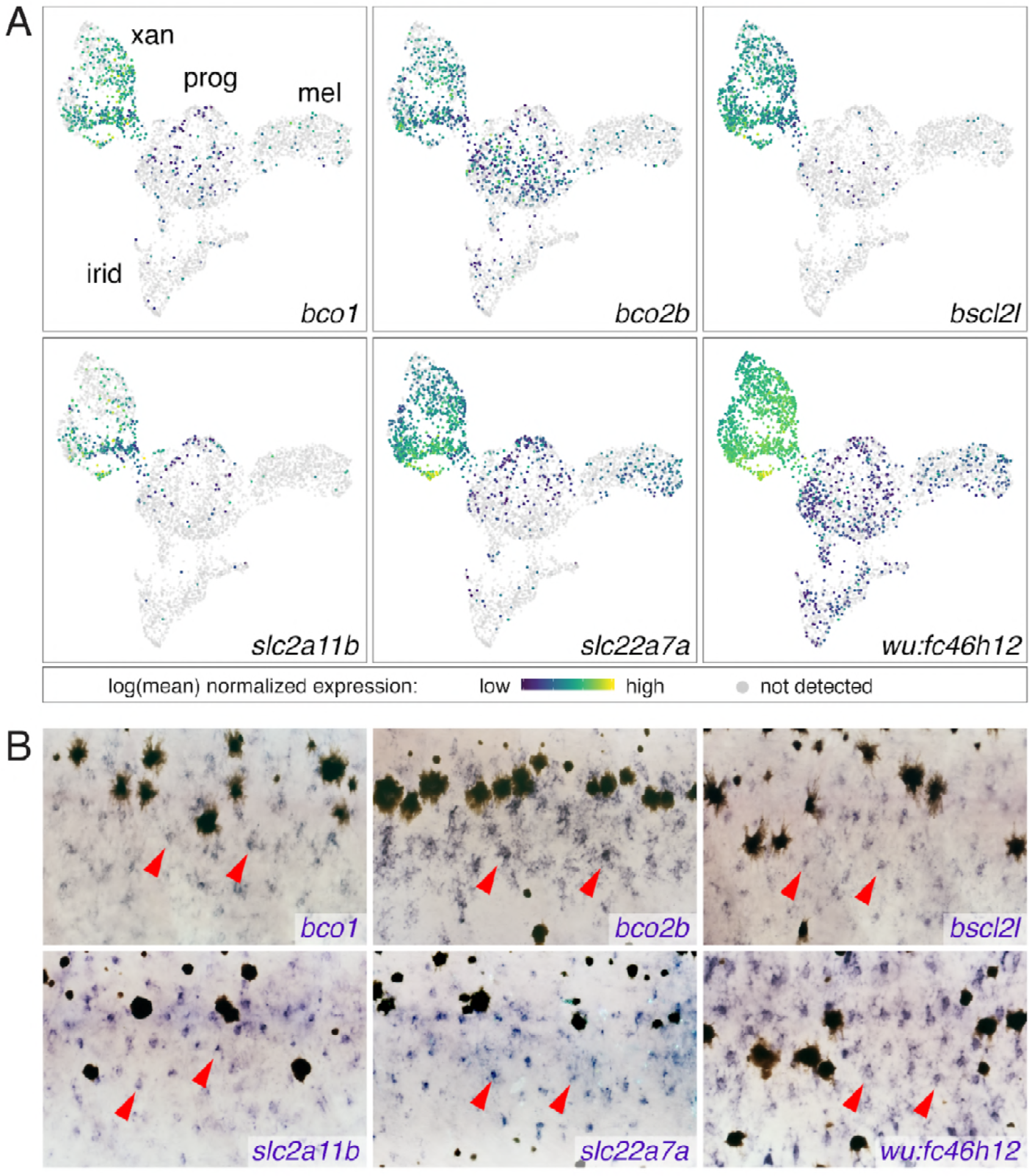
Xanthophore cluster-specific expression identifies novel xanthophore markers. **(A)** UMAP plots of pigment cells colored by expression of xanthophore cluster-enriched genes (*bco1, bco2b, bscl2l, slc2a11b, slc22a7a, wu:fc46h12*). **(B)** Expression in xanthophores (e.g., red arrowheads) of genes shown in A, as revealed by whole-mount *in situ* hybridization patterns corresponding to those of known xanthophore lineage markers and localization of differentiated and cryptic xanthophores (Hamada et al., 2014; Lang et al., 2009; McMenamin et al., 2014; Parichy et al., 2000b).

**Figure 3—figure supplement 3.**
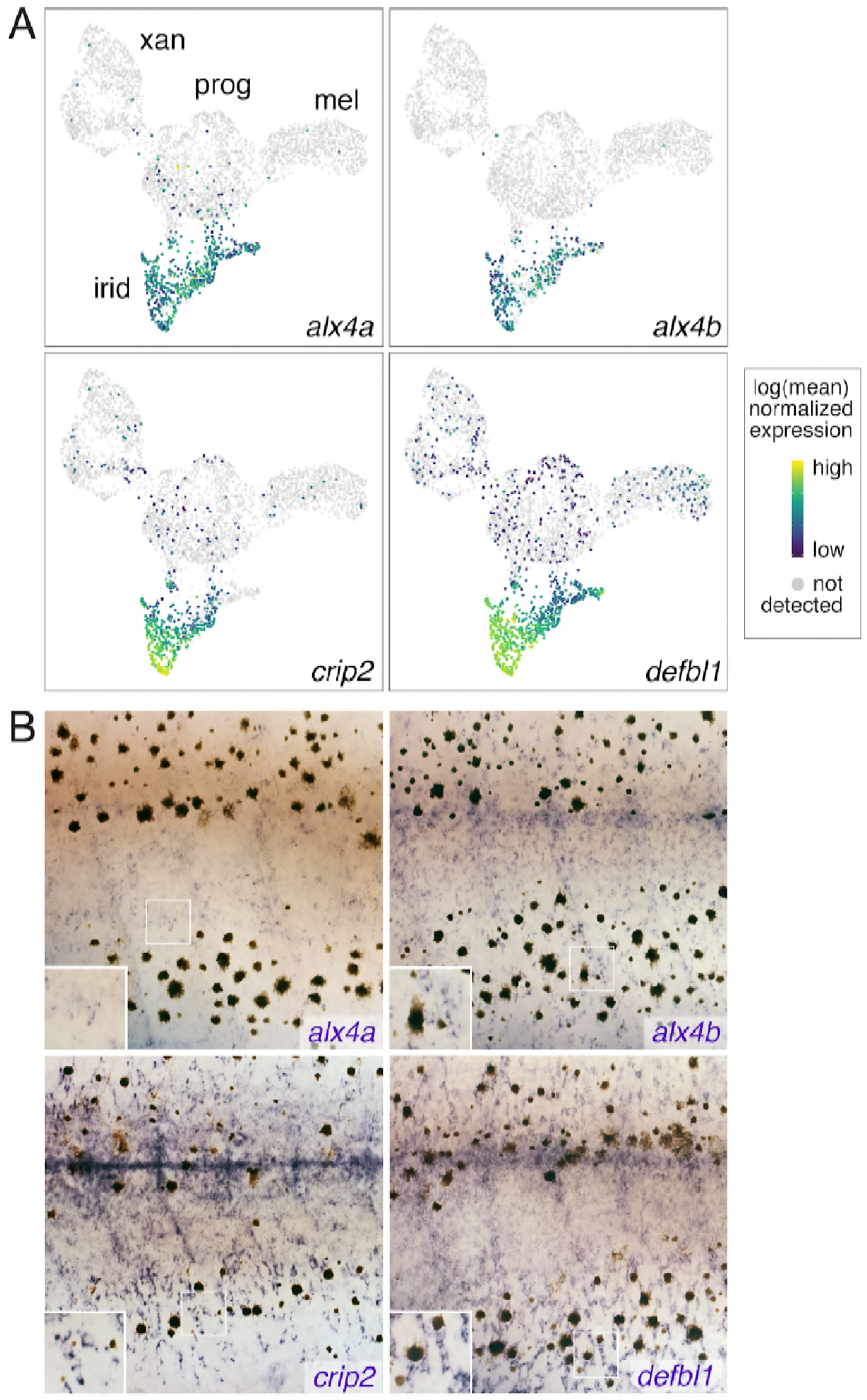
Iridophore cluster-specific expression identifies novel iridophore markers. **(A)** UMAP plots of pigment cells colored by expression of iridophore cluster-specific genes (*alx4a, alx4b, crip2, defbl1*). **(B)** Whole-mount in situ hybridization of genes in A reveals patterns corresponding to previously described iridophore markers and locations (Lang et al., 2009; Patterson and Parichy, 2013; Spiewak et al., 2018). Insets, higher magnification views of blue-stained iridophores.

**Figure 3—figure supplement 4.**
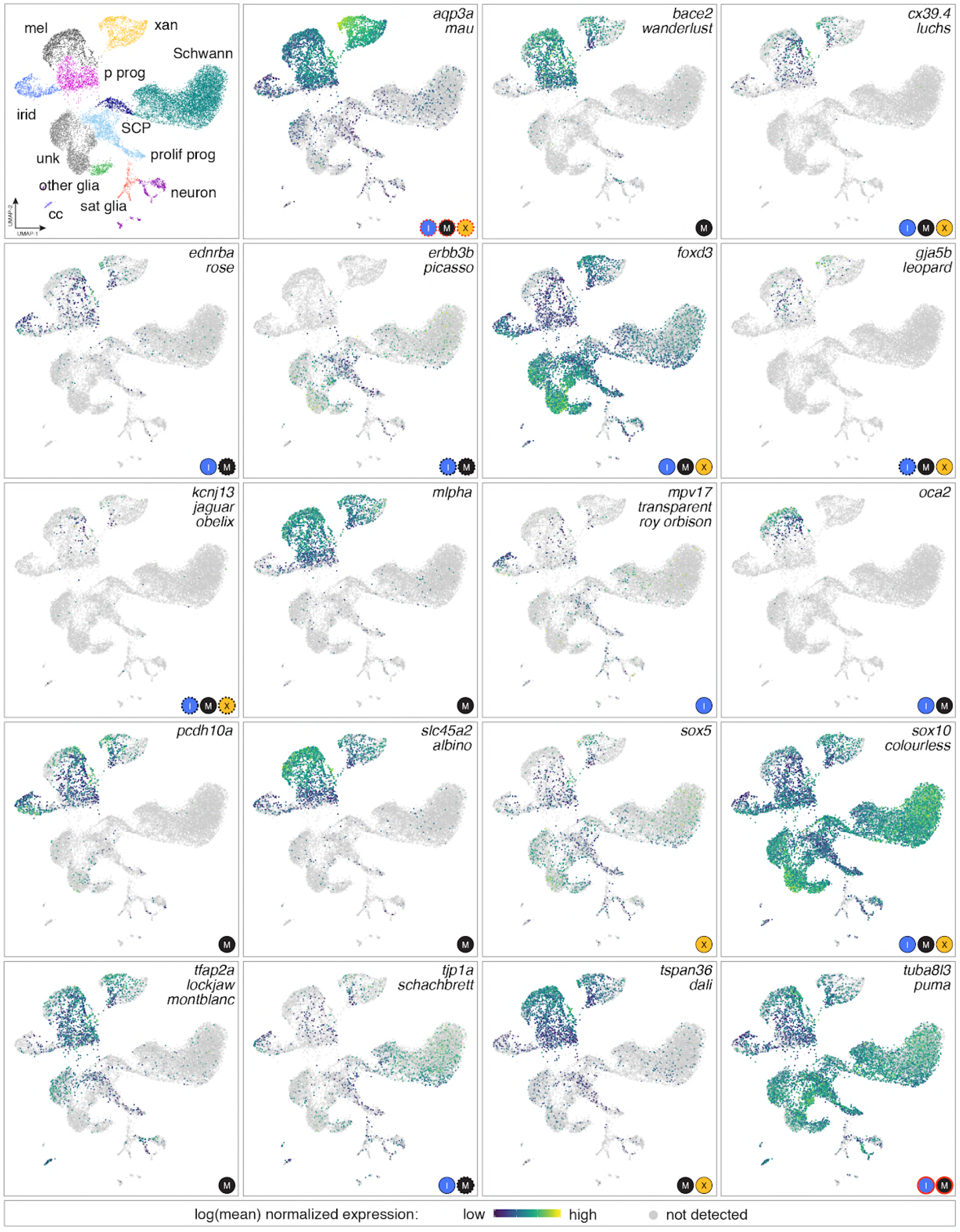
Genes identified as zebrafish pigmentation mutants often had expression domains beyond affected cell types. Mutations affecting a variety of pigmentary traits have been recovered or induced and affect pigment deposition, specification or morphogenesis of one or more pigment cell classes, and pattern at EL, adult or both stages (Arduini et al., 2009; Barrallo-Gimeno et al., 2004; Beirl et al., 2014; Budi et al., 2008; D’Agati et al., 2017; Dooley et al., 2013b; Dutton et al., 2001; Eskova et al., 2017; Fadeev et al., 2015; Inoue et al., 2014; Irion et al., 2014; Iwashita et al., 2006; Knight et al., 2003; Krauss et al., 2013; Larson et al., 2010; Nagao et al., 2018; Parichy et al., 2000a; Sheets et al., 2007; Watanabe et al., 2006; Williams et al., 2018; Zhang et al., 2018). Shown are expression domains observed for affected genes in scRNA-Seq analyses of adult NC-derived cells (euthyroid and hypothyroid). Gene names are listed at upper right of each box, with corresponding mutant names indicated below for those loci identified in forward genetic screens (for mutants isolated independently in different screens more than one name is indicated). Logos at lower right of each panel are cell types reported to be affected. In some instances only EL phenotypes have been reported (e.g., *pcdh10a*). Red outlines around cell type logos indicates neomorphic alleles (*aqp3a*, *tuba8l3a*); dashed lines indicate effects that are known to be non-autonomous to the affected cell types (e.g., *erbb3b*). In many instances scRNA-Seq expression domains identified a more diverse array of cell types than would be expected from gross mutant phenotypes alone. For example, *slc45a2* is required for melanization of melanophores but detected at lower levels in xanthophores and iridophores, whereas *mlpha* is required for melanosome dispersion but also expressed in xanthophores. Such instances raise the possibility of cell-type specific expression that is not functionally significant (e.g., if other pathway members are not themselves expression), unanticipated functions that result in only subtle loss-of-function phenotypes not yet identified, or amelioration of functional deficiencies by cell-type specific mechanisms of genetic compensation. Conversely, genes not expressed in affected cell types suggest or support prior inferences of non-autonomous functions, or requirements in a common progenitor (*erbb3b*, *oca2*). Genes for some well-studied mutants [e.g., *ltk*/*sparse (Johnson et al., 1995; Parichy et al., 1999)*] were expressed at levels too low to be detected by scRNA-Seq.

**Figure 3—figure supplement 5.**
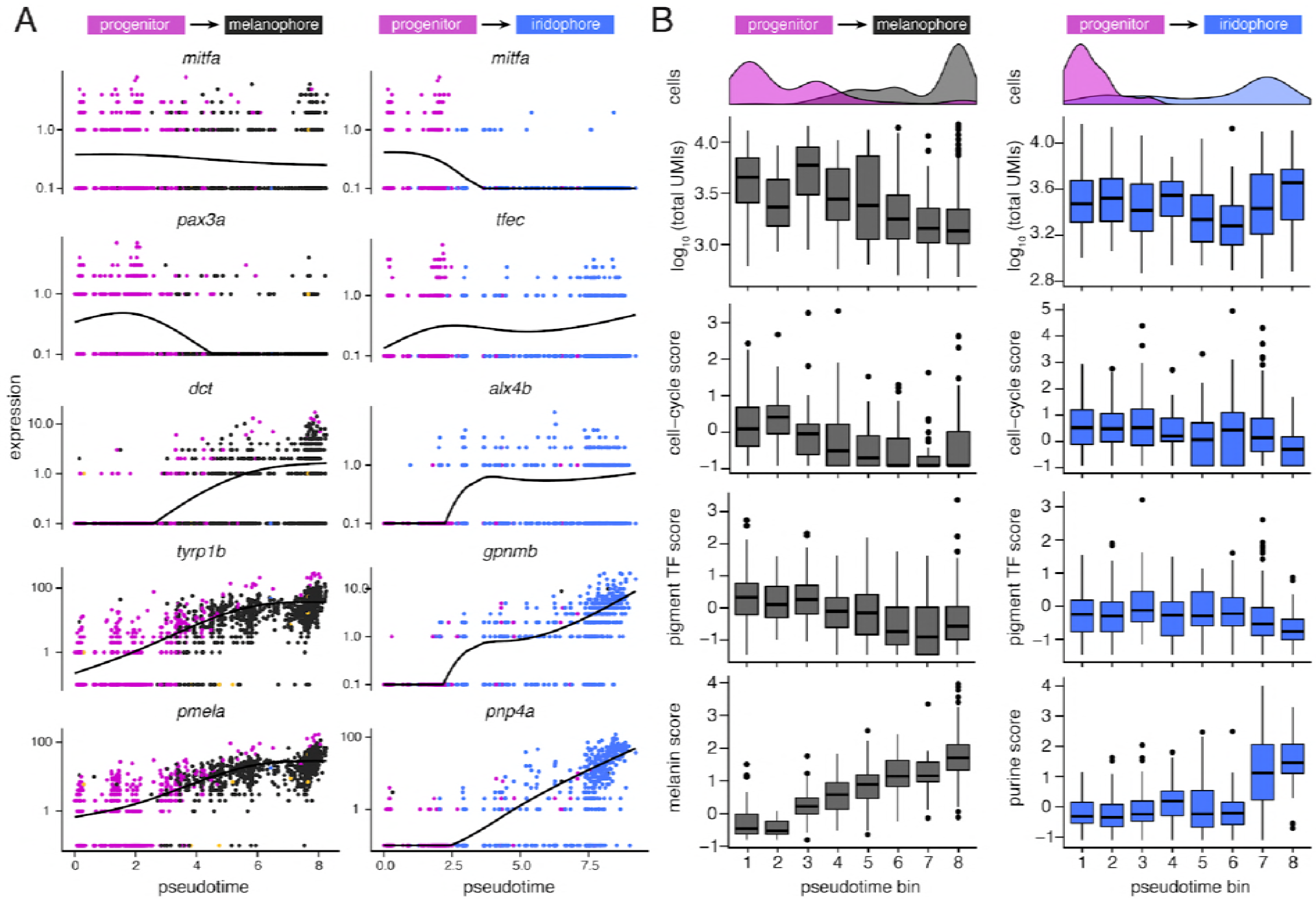
Dynamics of gene expression over pseudotime recapitulated distinct melanophore and iridophore differentiation programs. **(A)** Expression of genes over pseudotime reflect predicted kinetics for melanophores and iridophores. Solid lines indicate smoothed expression curves for all cells in the branch. *mitfa* expression declined only marginally with melanophore differentiation yet decreased markedly with a transition from progenitor to iridophore as expected(Curran et al., 2010). *pax3a* was expressed in pigment progenitors (magenta) and decreased across pseudotime in melanophores, whereas expression of *tfec*, a transcription factor expressed in iridophores(Lister et al., 2011), increased over pseudotime. Melanin synthesis enzyme genes, *dct* and *tryp1b*, as well as *pmel*, encoding a melanosome-associated transmembrane protein, all increased over pseudotime in melanophores. In iridophores, *gpnmb* and *pnp4a* showed elevated expression late in pseudotime, as expected (Curran et al., 2010; Higdon et al., 2013). **(B)** Trends of total transcript UMI counts, scores of expressed cell-cycle (e.g. *ccnd1, pcna*), pigment cell transcription factors (e.g. *mitfa, tfec, pax7b*, *tfap2a*), and pigment synthesis-related genes (e.g. *impdh1b, gart*, and *atic* for purine processing in iridophores; *tyrp1b, pmela*, and *tyr* for melanin synthesis in melanophores) in bins across pseudotime for melanophore and iridophore trajectory branches (for all score-associated genes, see **Table 4**). Histograms indicate cell-type specific densities across pseudotime for each branch. For melanophores, total transcript number per cell decreased over pseudotime and expression levels of melanin synthesis genes increased. In iridophores, mRNA levels stayed relatively constant whereas expression of purine synthesis genes increased. The expression score for cell-cycle genes was greater for iridophores than melanophores at the terminal step of pseudotime (*P*<0.0002; Wilcoxon; pseudotime bin 8; *n*=91 iridophores, 319 melanophores), consistent with iridophores continuing to proliferate even after differentiation, and melanophores normally failing to do so (Budi et al., 2011; Darzynkiewicz et al., 1980; McMenamin et al., 2014; Spiewak et al., 2018).

**Figure 5—figure supplement 1.**
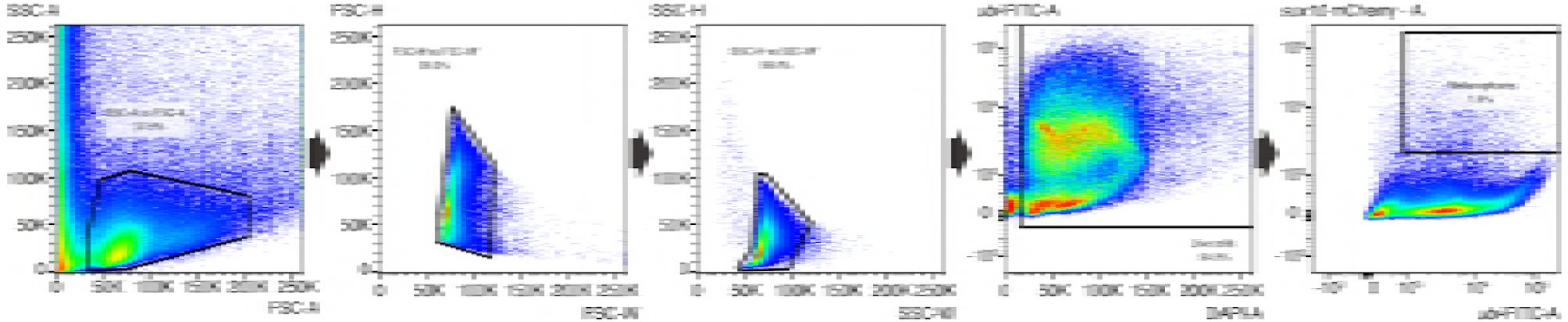
Melanophore FACS gating. Sequential FACS gating strategy (left to right) used in determining Lysotracker normalized MFI. For details, see Materials and Methods. (SSC, side scatter; FSC, forward scatter; W, width; H, height; A, area).

**Figure 6—figure supplement 1.**
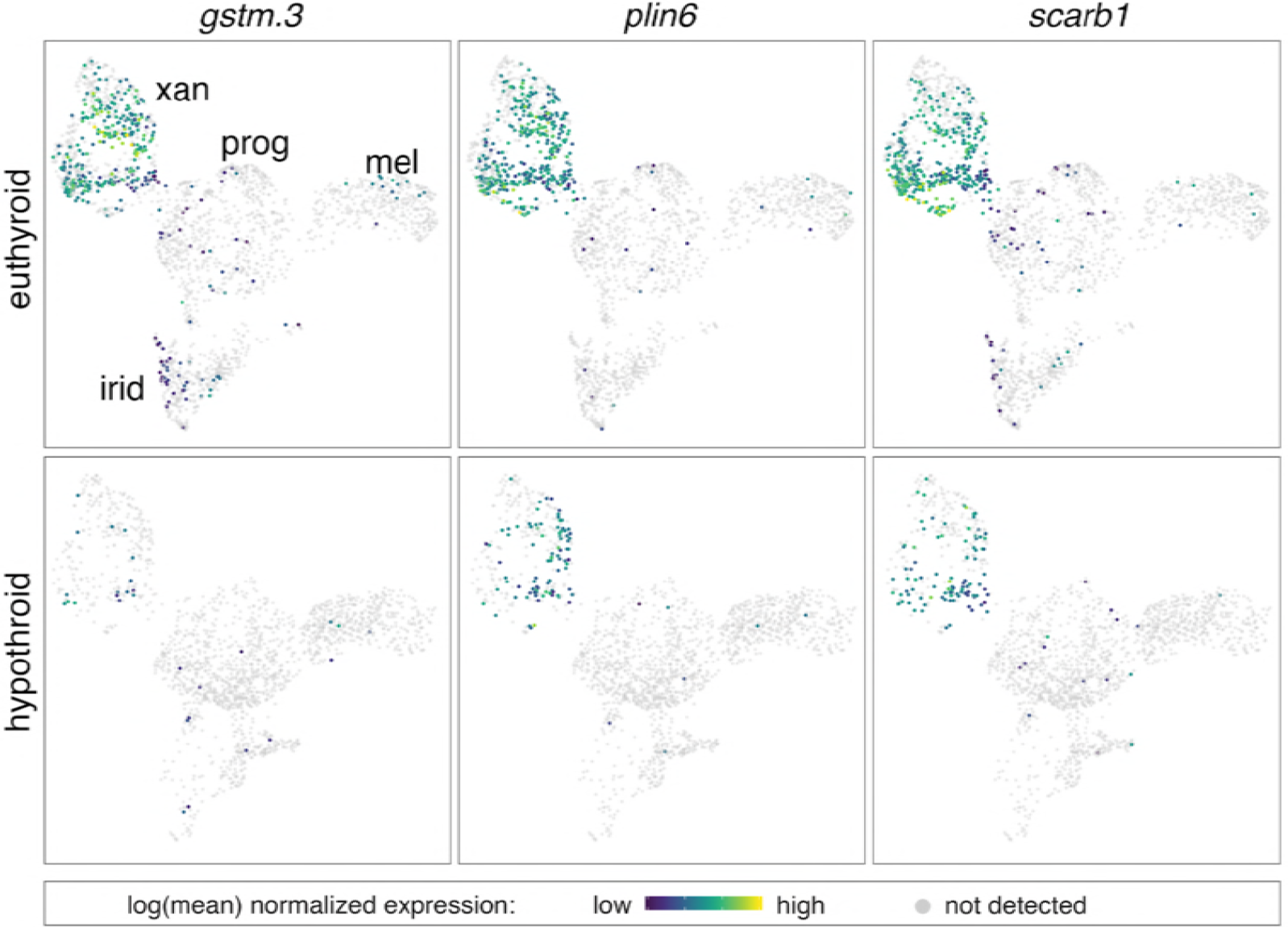
Expression of multiple carotenoid-related genes in xanthophores are affected by TH. UMAP plots of pigment cell clusters colored by expression of TH-dependent genes in xanthophores: *gstm.3* (*q*=6.9E-99, log_2_FC = 4.3)*, plin6* (*q*= 1.9E-13, log_2_FC=1.3)*, scarb1* (*q*=3.5E-11, log_2_FC=1.2).

**Figure 6—figure supplement 2.**
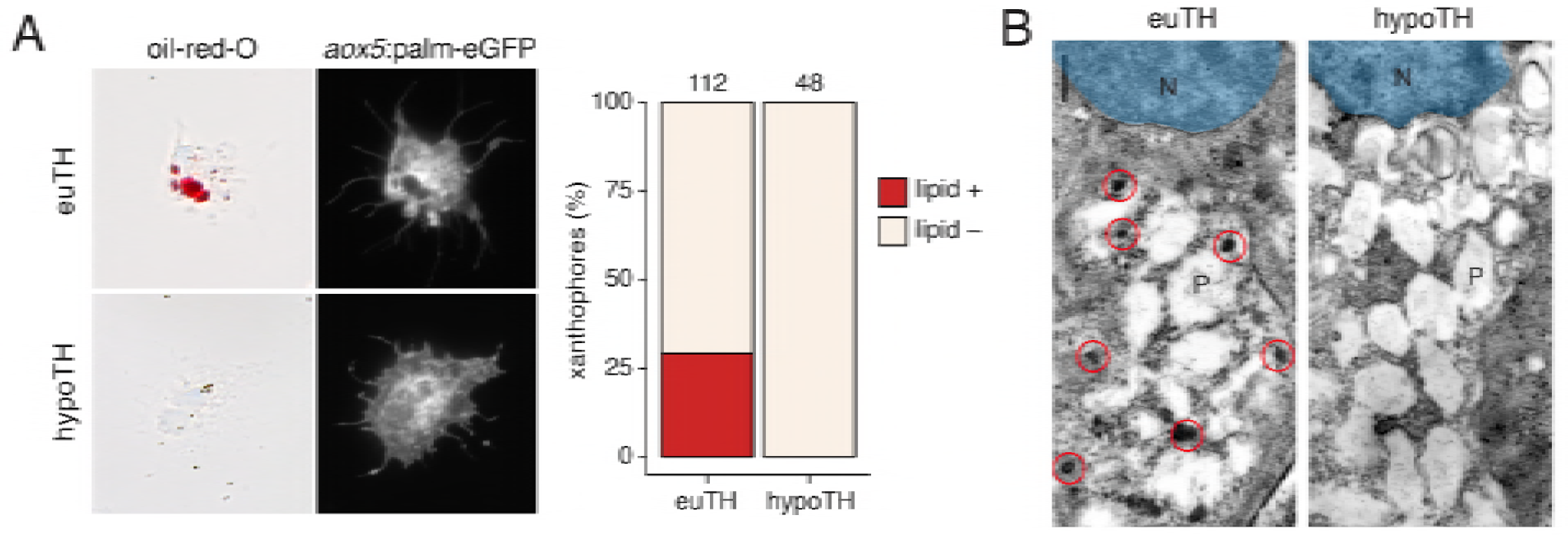
TH promotes development of lipid-filled carotenoid droplets in xanthophores. **(A)** Carotenoid pigments are normally localized to lipid droplets, the presence of which can be revealed by Oil-red-O staining. Here, a proportion of *aox5*:palmEGFP+ xanthophores stained *ex vivo* from euthyroid fish (*n*=112 cells) contained lipid (red), whereas xanthophores from hypothyroid fish (*n*=48 cells) were never observed to have such lipid contents. **(B)** Ultrastructurally, carotenoids and lipids are detectable as electron-dense carotenoid vesicles (red circles) (Djurdjevič et al., 2015; Granneman et al., 2017; Obika, 1993), which were observed in xanthophores from euthyroid but not hypothyroid fish. N, nucleus. P, pterinosome—the pteridine-containing organelle of xanthophores (Bagnara et al., 1968; Hirata et al., 2003; Matsumoto, 1965; Obika, 1993).

**Figure 6—figure supplement 3.**
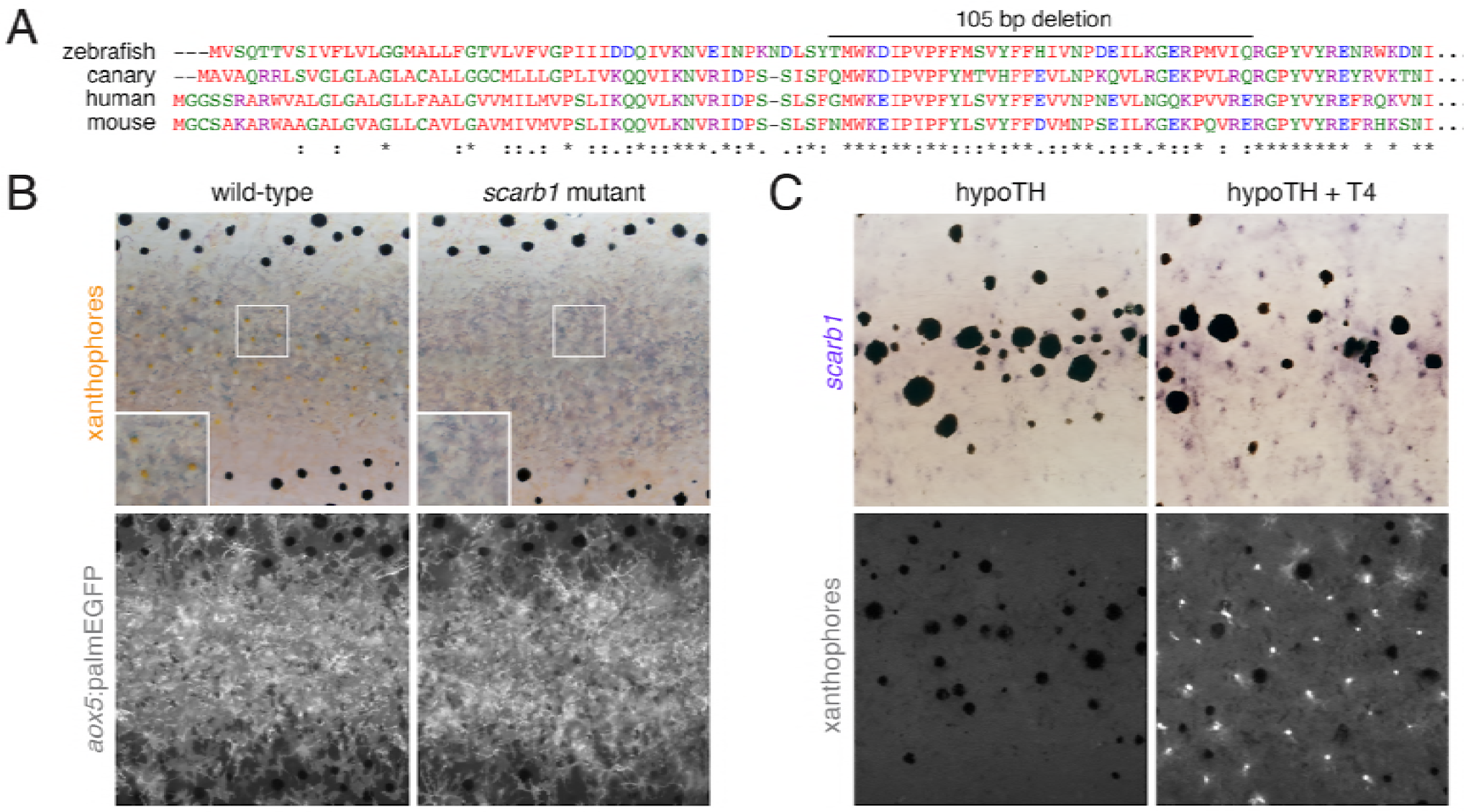
*scarb1* is specifically involved in xanthophore maturation and is induced by TH. **(A)** Scarb1 protein alignment. Zebrafish *scarb1* mutants had a 105 bp, in-frame deletion in a conserved region of the protein. **(B)** *scarb1* mutants lacked mature, interstripe xanthophores but had normal stripes and *aox5:palmEGFP* expression, suggesting that patterning and unpigmented xanthophores were normal. **(C)** In hypothyroid fish treated with exogenous TH (T4), *scarb1* expression was rescued within ~1 d (upper) and carotenoid autofluorescence of xanthophores was recovered within ~2 d (lower).

**Figure 6—figure supplement 4.**
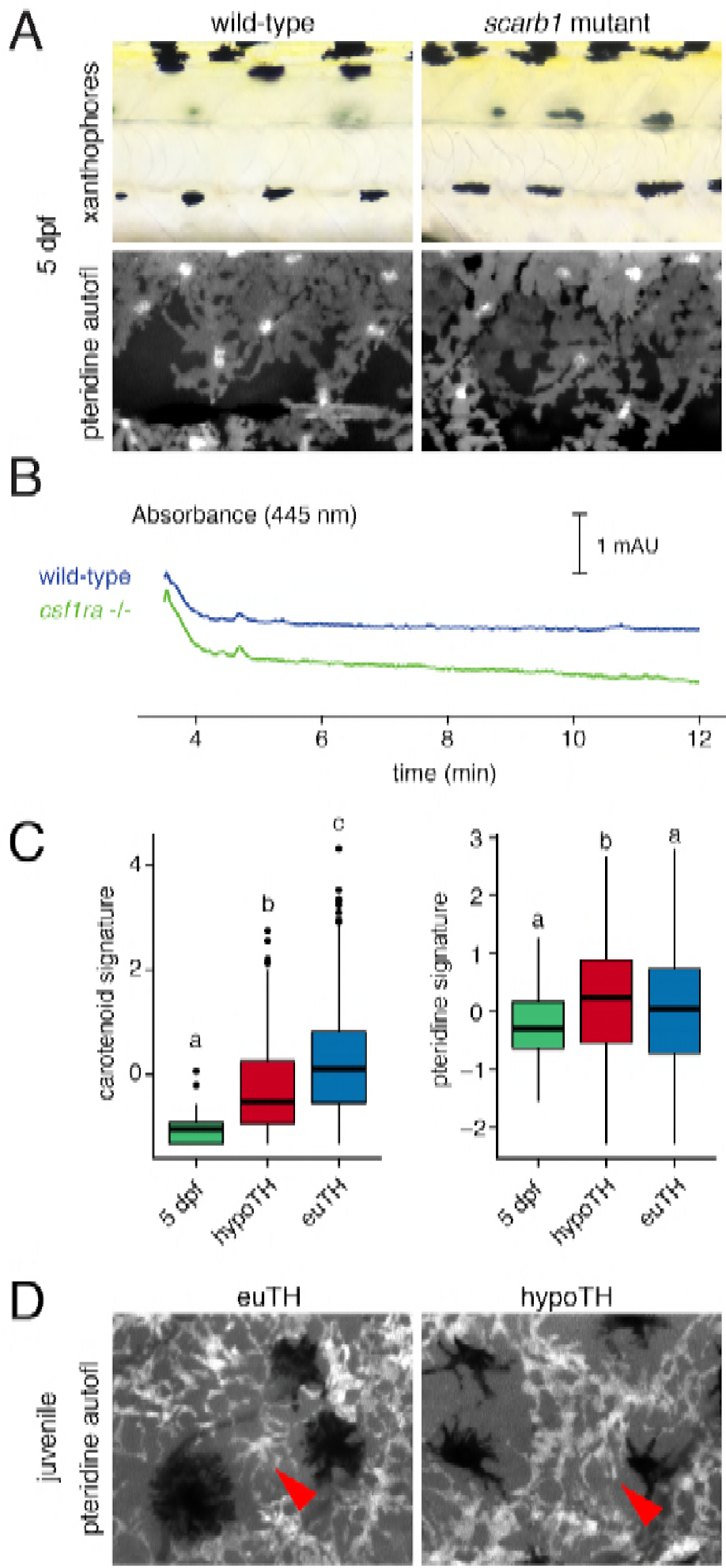
Xanthophores switch yellow pigmentation programs during the larval-to-adult transition. **(A)** At 5 dpf, *scarb1* mutants had yellow larval xanthophores with wild-type levels of pteridines. **(B)** Carotenoids were not detectable in EL zebrafish (5 dpf, wild-type; compare to Fig. 3d); *csf1ra* mutants, which lack xanthophores, had HPLC profiles indistinguishable from wild-type. **(C)** Carotenoid and pteridine pathway signature scores for xanthophores in euthyroid EL and euthyroid and hypothyroid adult scRNA-Seq data sets. Box plots as in Fig. 3 with different letters above data indicating significant differences in *post hoc* comparisons (carotenoid - *P*<2e-16, pteridine - *P*=0.01, Tukey HSD). Pteridine signatures between EL, hypothyroid, and euthyroid xanthophores were more similar than carotenoid signatures. **(D)** Ammonia-induced pteridine fluorescence was present in adult xanthophores of both euthyroid and hypothyroid fish (red arrowheads).

**Figure 7—figure supplement 1.**
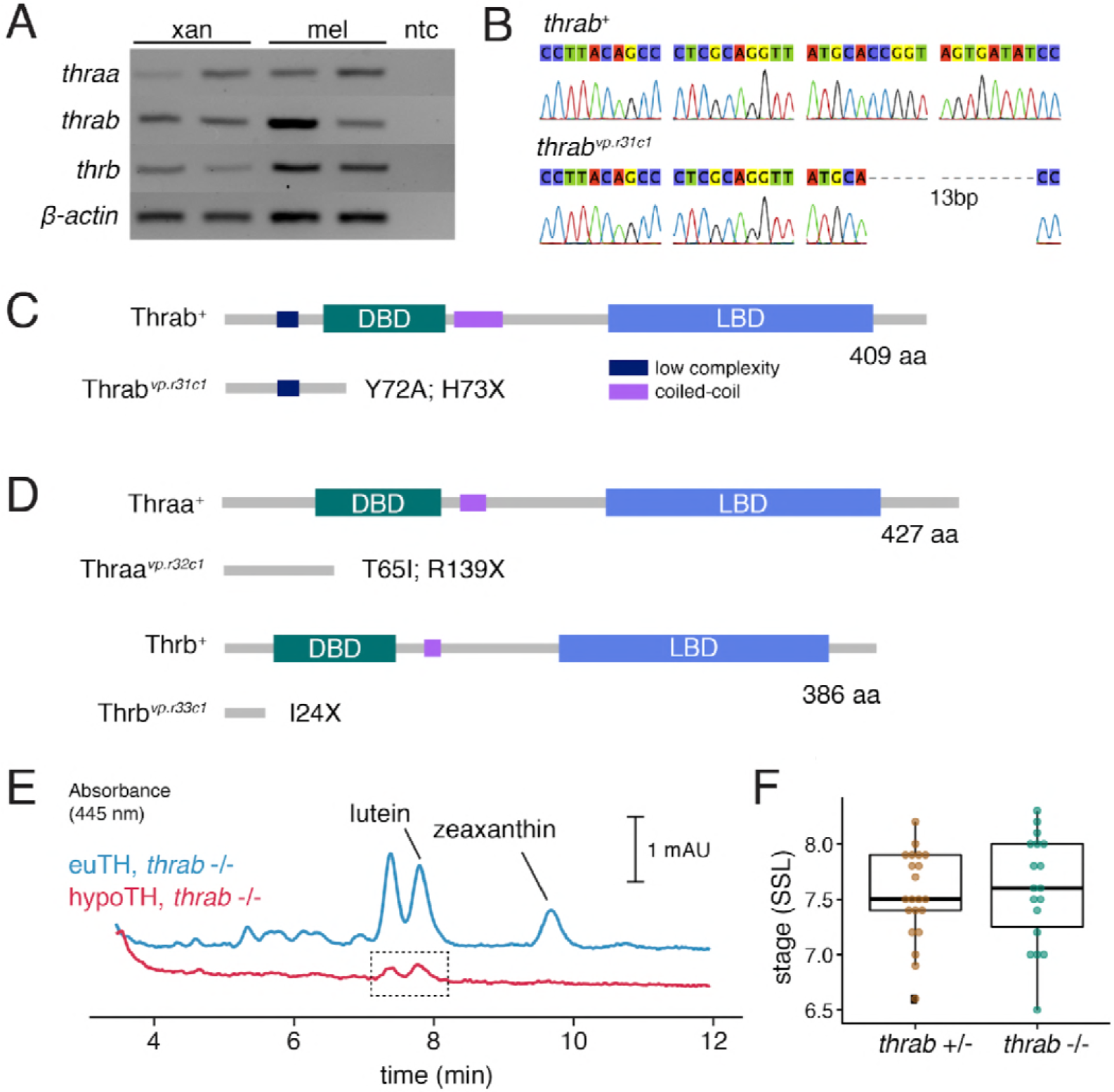
Zebrafish TR gene expression and mutants. **(A)** RT-PCR for *thraa, thrab*, and *thrb* in xanthophores and melanophores sorted by FACS for *aox5*:palmEGFP and *tyrp1b*:palm-mCherry, respectively. **(B)** Sanger sequencing of CRISPR/Cas9-induced mutant allele of *thrab* revealed a 13 bp deletion. **(C)** Schematic of Thrab wild-type and mutant proteins illustrating introduction of a novel amino acid followed by a premature stop codon at position 73. DBD, DNA binding domain; LBD, ligand binding domain. **(D)** Additional CRISPR/Cas9 mutant alleles for *thraa* and *thrb* had phenotypes indistinguishable from wild-type or *thrab* (**Figure 7A**). **(E)** HPLC revealed persisting carotenoids in hypothyroid fish mutant for *thrab* (boxed region), in contrast to the absence of detectable carotenoids in hypothyroid fish that were wild-type for *thrab* (**Figure 6D**). **(F)** Stage of first xanthophore appearance did not differ significantly (*P*=0.7) between euthyroid fish that were heterozygous or homozygous wild-type for *thrab* mutation.

## Other Supplementary Information includes

### Supplementary HTML

Interactive 3-dimensional UMAP representation of transcriptomic space. Cells are colored by type corresponding to **Figure 2A**.

**SaundersLM_tables.xlsx** containing:

Table 1. Genes enriched in specific cell types from post-embryonic NC derivatives.
Table 2. Genes from BEAM analysis of pigment cell lineages. Genes corresponding to rows in Fig. 2e by cluster.
Table 3. scRNA-seq sample information.
Table 4. Signature score genes.
Table 5. Ordering genes used as input for Monocle trajectory analysis.
Table 6. TH-dependent genes over pseudotime in melanophores. Genes correspond to clusters in extended data figure 12. Highlighted genes are have published roles in melanophores (with associated PMIDs).
Table 7. sgRNA and oligonucleotide sequences.

**Accompanying manuscript in revision** (Cao et al.)

## References

Adameyko I, Lallemend F, Aquino JB, Pereira JA, Topilko P, Müller T, Fritz N, Beljajeva A, Mochii M, Liste I, Usoskin D, Suter U, Birchmeier C, Ernfors P. 2009. Schwann cell precursors from nerve innervation are a cellular origin of melanocytes in skin. Cell 139:366–379.

Arduini BL, Bosse KM, Henion PD. 2009. Genetic ablation of neural crest cell diversification. Development 136:1987–1994.

Atchley WR, Hall BK. 1991. A model for development and evolution of complex morphological structures. Biol Rev Camb Philos Soc 66:101–157.

Bagnara JT, Matsumoto J. 2006. Comparative Anatomy and Physiology of Pigment Cells in Nonmammalian Tissues In: Nordlund JJ, Boissy RE, Hearing VJ, King RA, Oetting WS, Ortonne J-P, editors. The Pigmentary System. Oxford, UK: Blackwell Publishing Ltd. pp. 11–59.

Bagnara JT, Taylor JD, Hadley ME. 1968. The dermal chromatophore unit. J Cell Biol 38:67–79.

Barrallo-Gimeno A, Holzschuh J, Driever W, Knapik EW. 2004. Neural crest survival and differentiation in zebrafish depends on mont blanc/tfap2a gene function. Development 131:1463–1477.

Baxter LL, Watkins-Chow DE, Pavan WJ, Loftus SK. 2018. A curated gene list for expanding the horizons of pigmentation biology. Pigment Cell Melanoma Res. doi:10.1111/pcmr.12743

Becht E, McInnes L, Healy J, Dutertre C-A, Kwok IWH, Ng LG, Ginhoux F, Newell EW. 2018. Dimensionality reduction for visualizing single-cell data using UMAP. Nat Biotechnol. doi:10.1038/nbt.4314

Beirl AJ, Linbo TH, Cobb MJ, Cooper CD. 2014. oca2 Regulation of chromatophore differentiation and number is cell type specific in zebrafish. Pigment Cell Melanoma Res 27:178–189.

Blondel VD, Guillaume J-L, Lambiotte R, Lefebvre E. 2008. Fast unfolding of communities in large networks. arXiv [physics.soc-ph].

Braasch I, Brunet F, Volff J-N, Schartl M. 2009. Pigmentation pathway evolution after whole-genome duplication in fish. Genome Biol Evol 1:479–493.

Braasch I, Peterson SM, Desvignes T, McCluskey BM, Batzel P, Postlethwait JH. 2015. A new model army: Emerging fish models to study the genomics of vertebrate Evo-Devo. J Exp Zool B Mol Dev Evol 324:316–341.

Brent GA. 2012. Mechanisms of thyroid hormone action. J Clin Invest 122:3035–3043.

Brown DD, Cai L. 2007. Amphibian metamorphosis. Dev Biol 306:20–33.

Buchholz DR, Hsia S-CV, Fu L, Shi Y-B. 2003. A dominant-negative thyroid hormone receptor blocks amphibian metamorphosis by retaining corepressors at target genes. Mol Cell Biol 23:6750–6758.

Budi EH, Patterson LB, Parichy DM. 2011. Post-embryonic nerve-associated precursors to adult pigment cells: genetic requirements and dynamics of morphogenesis and differentiation. PLoS Genet 7:e1002044.

Budi EH, Patterson LB, Parichy DM. 2008. Embryonic requirements for ErbB signaling in neural crest development and adult pigment pattern formation. Development 135:2603–2614.

Cao J, Spielmann M, Qiu X, Huang X, Ibrahim DM, Hill AJ, Zhang F, Mundlos S, Christiansen L, Steemers FJ, Trapnell C, Shendure J. 2018. The dynamic transcriptional landscape of mammalian organogenesis at single cell resolution. Submitted.

Ceol CJ, Houvras Y, Jane-Valbuena J, Bilodeau S, Orlando DA, Battisti V, Fritsch L, Lin WM, Hollmann TJ, Ferré F, Bourque C, Burke CJ, Turner L, Uong A, Johnson LA, Beroukhim R, Mermel CH, Loda M, Ait-Si-Ali S, Garraway LA, Young RA, Zon LI. 2011. The histone methyltransferase SETDB1 is recurrently amplified in melanoma and accelerates its onset. Nature 471:513–517.

Chang J, Wang M, Gui W, Zhao Y, Yu L, Zhu G. 2012. Changes in thyroid hormone levels during zebrafish development. Zoolog Sci 29:181–184.

Chinenov Y, Kerppola TK. 2001. Close encounters of many kinds: Fos-Jun interactions that mediate transcription regulatory specificity. Oncogene 20:2438–2452.

Choi J, Suzuki K-IT, Sakuma T, Shewade L, Yamamoto T, Buchholz DR. 2015. Unliganded thyroid hormone receptor α regulates developmental timing via gene repression in Xenopus tropicalis. Endocrinology 156:735–744.

Coller HA, Sang L, Roberts JM. 2006. A new description of cellular quiescence. PLoS Biol 4:e83.

Curran K, Lister JA, Kunkel GR, Prendergast A, Parichy DM, Raible DW. 2010. Interplay between Foxd3 and Mitf regulates cell fate plasticity in the zebrafish neural crest. Dev Biol 344:107–118.

D’Agati G, Beltre R, Sessa A, Burger A, Zhou Y, Mosimann C, White RM. 2017. A defect in the mitochondrial protein Mpv17 underlies the transparent casper zebrafish. Dev Biol 430:11–17.

Darzynkiewicz Z, Traganos F, Melamed MR. 1980. New cell cycle compartments identified by multiparameter flow cytometry. Cytometry 1:98–108.

Dimri GP, Lee X, Basile G, Acosta M, Scott G, Roskelley C, Medrano EE, Linskens M, Rubelj I, Pereira-Smith O. 1995. A biomarker that identifies senescent human cells in culture and in aging skin in vivo. Proc Natl Acad Sci U S A 92:9363–9367.

Djurdjevič I, Kreft ME, Sušnik Bajec S. 2015. Comparison of pigment cell ultrastructure and organisation in the dermis of marble trout and brown trout, and first description of erythrophore ultrastructure in salmonids. J Anat 227:583–595.

Dooley CM, Mongera A, Walderich B, Nüsslein-Volhard C. 2013a. On the embryonic origin of adult melanophores: the role of ErbB and Kit signalling in establishing melanophore stem cells in zebrafish. Development 140:1003–1013.

Dooley CM, Schwarz H, Mueller KP, Mongera A, Konantz M, Neuhauss SCF, Nüsslein-Volhard C, Geisler R. 2013b. Slc45a2 and V-ATPase are regulators of melanosomal pH homeostasis in zebrafish, providing a mechanism for human pigment evolution and disease. Pigment Cell Melanoma Res 26:205–217.

Dutton KA, Pauliny A, Lopes SS, Elworthy S, Carney TJ, Rauch J, Geisler R, Haffter P, Kelsh RN. 2001. Zebrafish colourless encodes sox10 and specifies non-ectomesenchymal neural crest fates. Development 128:4113–4125.

Ebisuya M, Briscoe J. 2018. What does time mean in development? Development 145. doi:10.1242/dev.164368

Ellerhorst JA, Cooksley CD, Broemeling L, Johnson MM, Grimm EA. 2003. High prevalence of hypothyroidism among patients with cutaneous melanoma. Oncol Rep 10:1317–1320.

Eom DS, Bain EJ, Patterson LB, Grout ME, Parichy DM. 2015. Long-distance communication by specialized cellular projections during pigment pattern development and evolution. Elife 4. doi:10.7554/eLife.12401

Eskova A, Chauvigné F, Maischein H-M, Ammelburg M, Cerdà J, Nüsslein-Volhard C, Irion U. 2017. Gain-of-function mutations in Aqp3a influence zebrafish pigment pattern formation through the tissue environment. Development 144:2059–2069.

Fadeev A, Krauss J, Frohnhöfer HG, Irion U, Nüsslein-Volhard C. 2015. Tight Junction Protein 1a regulates pigment cell organisation during zebrafish colour patterning. Elife 4. doi:10.7554/eLife.06545

Flamant F, Poguet A-L, Plateroti M, Chassande O, Gauthier K, Streichenberger N, Mansouri A, Samarut J. 2002. Congenital Hypothyroid Pax8−/− Mutant Mice Can Be Rescued by Inactivating the TRα Gene. Mol Endocrinol 16:24–32.

Flamant F, Samarut J. 2003. Thyroid hormone receptors: lessons from knockout and knock-in mutant mice. Trends Endocrinol Metab 14:85–90.

Frohnhöfer HG, Krauss J, Maischein H-M, Nüsslein-Volhard C. 2013. Iridophores and their interactions with other chromatophores are required for stripe formation in zebrafish. Development 140:2997–3007.

Gans C, Northcutt RG. 1983. Neural crest and the origin of vertebrates: a new head. Science 220:268–273.

Granneman JG, Kimler VA, Zhang H, Ye X, Luo X, Postlethwait JH, Thummel R. 2017. Lipid droplet biology and evolution illuminated by the characterization of a novel perilipin in teleost fish. Elife 6. doi:10.7554/eLife.21771

Hamada H, Watanabe M, Lau HE, Nishida T, Hasegawa T, Parichy DM, Kondo S. 2014. Involvement of Delta/Notch signaling in zebrafish adult pigment stripe patterning. Development 141:318–324.

Higdon CW, Mitra RD, Johnson SL. 2013. Gene expression analysis of zebrafish melanocytes, iridophores, and retinal pigmented epithelium reveals indicators of biological function and developmental origin. PLoS One 8:e67801.

Hirata M, Nakamura K-I, Kanemaru T, Shibata Y, Kondo S. 2003. Pigment cell organization in the hypodermis of zebrafish. Dev Dyn 227:497–503.

Hörlein AJ, Näär AM, Heinzel T, Torchia J, Gloss B, Kurokawa R, Ryan A, Kamei Y, Söderström M, Glass CK. 1995. Ligand-independent repression by the thyroid hormone receptor mediated by a nuclear receptor co-repressor. Nature 377:397–404.

Hultman KA, Bahary N, Zon LI, Johnson SL. 2007. Gene Duplication of the zebrafish kit ligand and partitioning of melanocyte development functions to kit ligand a. PLoS Genet 3:e17.

Ilicic T, Kim JK, Kolodziejczyk AA, Bagger FO, McCarthy DJ, Marioni JC, Teichmann SA. 2016. Classification of low quality cells from single-cell RNA-seq data. Genome Biol 17:29.

Inoue S, Kondo S, Parichy DM, Watanabe M. 2014. Tetraspanin 3c requirement for pigment cell interactions and boundary formation in zebrafish adult pigment stripes. Pigment Cell Melanoma Res 27:190–200.

Irion U, Frohnhöfer HG, Krauss J, Çolak Champollion T, Maischein H-M, Geiger-Rudolph S, Weiler C, Nüsslein-Volhard C. 2014. Gap junctions composed of connexins 41.8 and 39.4 are essential for colour pattern formation in zebrafish. Elife 3:e05125.

Irion U, Singh AP, Nüsslein-Volhard C. 2016. The Developmental Genetics of Vertebrate Color Pattern Formation: Lessons from Zebrafish. Curr Top Dev Biol 117:141–169.

Iwashita M, Watanabe M, Ishii M, Chen T, Johnson SL, Kurachi Y, Okada N, Kondo S. 2006. Pigment pattern in jaguar/obelix zebrafish is caused by a Kir7.1 mutation: implications for the regulation of melanosome movement. PLoS Genet 2:e197.

Johnson SL, Africa D, Walker C, Weston JA. 1995. Genetic control of adult pigment stripe development in zebrafish. Dev Biol 167:27–33.

Kague E, Gallagher M, Burke S, Parsons M, Franz-Odendaal T, Fisher S. 2012. Skeletogenic fate of zebrafish cranial and trunk neural crest. PLoS One 7:e47394.

Kaufman CK, Mosimann C, Fan ZP, Yang S, Thomas AJ, Ablain J, Tan JL, Fogley RD, van Rooijen E, Hagedorn EJ, Ciarlo C, White RM, Matos DA, Puller A-C, Santoriello C, Liao EC, Young RA, Zon LI. 2016. A zebrafish melanoma model reveals emergence of neural crest identity during melanoma initiation. Science 351:aad2197.

Kelsh RN, Dutton K, Medlin J, Eisen JS. 2000a. Expression of zebrafish fkd6 in neural crest-derived glia. Mech Dev 93:161–164.

Kelsh RN, Schmid B, Eisen JS. 2000b. Genetic analysis of melanophore development in zebrafish embryos. Dev Biol 225:277–293.

Kelsh RN, Sosa KC, Owen JP, Yates CA. 2017. Zebrafish adult pigment stem cells are multipotent and form pigment cells by a progressive fate restriction process: Clonal analysis identifies shared origin of all pigment cell types. Bioessays 39. doi:10.1002/bies.201600234

Kiefer C, Sumser E, Wernet MF, Von Lintig J. 2002. A class B scavenger receptor mediates the cellular uptake of carotenoids in Drosophila. Proc Natl Acad Sci U S A 99:10581–10586.

Knight RD, Nair S, Nelson SS, Afshar A, Javidan Y, Geisler R, Rauch G-J, Schilling TF. 2003. lockjaw encodes a zebrafish tfap2a required for early neural crest development. Development 130:5755–5768.

Koopman R, Schaart G, Hesselink MK. 2001. Optimisation of oil red O staining permits combination with immunofluorescence and automated quantification of lipids. Histochem Cell Biol 116:63–68.

Krauss J, Astrinidis P, Frohnhöfer HG, Walderich B, Nüsslein-Volhard C. 2013. transparent, a gene affecting stripe formation in Zebrafish, encodes the mitochondrial protein Mpv17 that is required for iridophore survival. Biol Open 2:703–710.

Kurz DJ, Decary S, Hong Y, Erusalimsky JD. 2000. Senescence-associated (beta)-galactosidase reflects an increase in lysosomal mass during replicative ageing of human endothelial cells. J Cell Sci 113 (Pt 20):3613–3622.

Lang MR, Patterson LB, Gordon TN, Johnson SL, Parichy DM. 2009. Basonuclin-2 requirements for zebrafish adult pigment pattern development and female fertility. PLoS Genet 5:e1000744.

Larson TA, Gordon TN, Lau HE, Parichy DM. 2010. Defective adult oligodendrocyte and Schwann cell development, pigment pattern, and craniofacial morphology in puma mutant zebrafish having an alpha tubulin mutation. Dev Biol 346:296–309.

Lee BY, Han JA, Im JS, Morrone A, Johung K, Goodwin EC, Kleijer WJ, DiMaio D, Hwang ES. 2006. Senescence-associated beta-galactosidase is lysosomal beta-galactosidase. Aging Cell 5:187–195.

Lister JA, Lane BM, Nguyen A, Lunney K. 2011. Embryonic expression of zebrafish MiT family genes tfe3b, tfeb, and tfec. Dev Dyn 240:2529–2538.

Lister JA, Robertson CP, Lepage T, Johnson SL, Raible DW. 1999. nacre encodes a zebrafish microphthalmia-related protein that regulates neural-crest-derived pigment cell fate. Development 126:3757–3767.

Luo R, An M, Arduini BL, Henion PD. 2001. Specific pan-neural crest expression of zebrafish Crestin throughout embryonic development. Dev Dyn 220:169–174.

Mahalwar P, Walderich B, Singh AP, Nüsslein-Volhard C. 2014. Local reorganization of xanthophores fine-tunes and colors the striped pattern of zebrafish. Science 345:1362–1364.

Matsumoto J. 1965. Studies on fine structure and cytochemical properties of erythrophores in swordtail, Xiphophorus helleri, with special reference to their pigment granules (Pterinosomes). J Cell Biol 27:493–504.

McInnes L, Healy J, Saul N, Großberger L. 2018. UMAP: Uniform Manifold Approximation and Projection. JOSS 3:861.

McMenamin SK, Bain EJ, McCann AE, Patterson LB, Eom DS, Waller ZP, Hamill JC, Kuhlman JA, Eisen JS, Parichy DM. 2014. Thyroid hormone-dependent adult pigment cell lineage and pattern in zebrafish. Science 345:1358–1361.

Minchin JEN, Hughes SM. 2008. Sequential actions of Pax3 and Pax7 drive xanthophore development in zebrafish neural crest. Dev Biol 317:508–522.

Mosimann C, Kaufman CK, Li P, Pugach EK, Tamplin OJ, Zon LI. 2011. Ubiquitous transgene expression and Cre-based recombination driven by the ubiquitin promoter in zebrafish. Development 138:169–177.

Nagao Y, Takada H, Miyadai M, Adachi T, Seki R, Kamei Y, Hara I, Taniguchi Y, Naruse K, Hibi M, Kelsh RN, Hashimoto H. 2018. Distinct interactions of Sox5 and Sox10 in fate specification of pigment cells in medaka and zebrafish. PLoS Genet 14:e1007260.

Nord H, Dennhag N, Muck J, von Hofsten J. 2016. Pax7 is required for establishment of the xanthophore lineage in zebrafish embryos. Mol Biol Cell 27:1853–1862.

Obika M. 1993. Formation of pterinosomes and carotenoid granules in xanthophores of the teleost Oryzias latipes as revealed by the rapid-freezing and freeze-substitution method. Cell Tissue Res 271:81–86.

Odenthal J, Rossnagel K, Haffter P, Kelsh RN, Vogelsang E, Brand M, van Eeden FJ, Furutani-Seiki M, Granato M, Hammerschmidt M, Heisenberg CP, Jiang YJ, Kane DA, Mullins MC, Nüsslein-Volhard C. 1996. Mutations affecting xanthophore pigmentation in the zebrafish, Danio rerio. Development 123:391–398.

Orr-Weaver TL. 2015. When bigger is better: the role of polyploidy in organogenesis. Trends Genet 31:307–315.

Parichy DM, Elizondo MR, Mills MG, Gordon TN, Engeszer RE. 2009. Normal table of postembryonic zebrafish development: staging by externally visible anatomy of the living fish. Dev Dyn 238:2975–3015.

Parichy DM, Mellgren EM, Rawls JF, Lopes SS, Kelsh RN, Johnson SL. 2000a. Mutational analysis of endothelin receptor b1 (rose) during neural crest and pigment pattern development in the zebrafish Danio rerio. Dev Biol 227:294–306.

Parichy DM, Ransom DG, Paw B, Zon LI, Johnson SL. 2000b. An orthologue of the kit-related gene fms is required for development of neural crest-derived xanthophores and a subpopulation of adult melanocytes in the zebrafish, Danio rerio. Development 127:3031–3044.

Parichy DM, Rawls JF, Pratt SJ, Whitfield TT, Johnson SL. 1999. Zebrafish sparse corresponds to an orthologue of c-kit and is required for the morphogenesis of a subpopulation of melanocytes, but is not essential for hematopoiesis or primordial germ cell development. Development 126:3425–3436.

Parichy DM, Spiewak JE. 2015. Origins of adult pigmentation: diversity in pigment stem cell lineages and implications for pattern evolution. Pigment Cell Melanoma Res 28:31–50.

Patil C, Walter P. 2001. Intracellular signaling from the endoplasmic reticulum to the nucleus: the unfolded protein response in yeast and mammals. Curr Opin Cell Biol 13:349–355.

Patterson LB, Bain EJ, Parichy DM. 2014. Pigment cell interactions and differential xanthophore recruitment underlying zebrafish stripe reiteration and Danio pattern evolution. Nat Commun 5:5299.

Patterson LB, Parichy DM. 2013. Interactions with iridophores and the tissue environment required for patterning melanophores and xanthophores during zebrafish adult pigment stripe formation. PLoS Genet 9:e1003561.

Qiu X, Hill A, Packer J, Lin D, Ma Y-A, Trapnell C. 2017a. Single-cell mRNA quantification and differential analysis with Census. Nat Methods 14:309–315.

Qiu X, Mao Q, Tang Y, Wang L, Chawla R, Pliner HA, Trapnell C. 2017b. Reversed graph embedding resolves complex single-cell trajectories. Nat Methods 14:979–982.

Quigley IK, Turner JM, Nuckels RJ, Manuel JL, Budi EH, MacDonald EL, Parichy DM. 2004. Pigment pattern evolution by differential deployment of neural crest and post-embryonic melanophore lineages in Danio fishes. Development 131:6053–6069.

R Core Team. 2017. R: A Language and Environment for Statistical Computing.

Riabowol K, Schiff J, Gilman MZ. 1992. Transcription factor AP-1 activity is required for initiation of DNA synthesis and is lost during cellular aging. Proc Natl Acad Sci U S A 89:157–161.

Shah AN, Davey CF, Whitebirch AC, Miller AC, Moens CB. 2015. Rapid reverse genetic screening using CRISPR in zebrafish. Nat Methods 12:535–540.

Shah M, Orengo IF, Rosen T. 2006. High prevalence of hypothyroidism in male patients with cutaneous melanoma. Dermatol Online J 12:1.

Sharan SK, Thomason LC, Kuznetsov SG, Court DL. 2009. Recombineering: a homologous recombination-based method of genetic engineering. Nat Protoc 4:206–223.

Sheets L, Ransom DG, Mellgren EM, Johnson SL, Schnapp BJ. 2007. Zebrafish melanophilin facilitates melanosome dispersion by regulating dynein. Curr Biol 17:1721–1734.

Shi Y-B. 2013. Chapter Ten - Unliganded Thyroid Hormone Receptor Regulates Metamorphic Timing via the Recruitment of Histone Deacetylase Complexes In: Rougvie AE, O’Connor MB, editors. Current Topics in Developmental Biology. Academic Press. pp. 275–297.

Shi Y-B. 1999. Amphibian Metamorphosis: From Morphology to Molecular Biology. Wiley-Liss.

Singh AP, Dinwiddie A, Mahalwar P, Schach U, Linker C, Irion U, Nüsslein-Volhard C. 2016. Pigment Cell Progenitors in Zebrafish Remain Multipotent through Metamorphosis. Dev Cell 38:316–330.

Singh AP, Schach U, Nüsslein-Volhard C. 2014. Proliferation, dispersal and patterned aggregation of iridophores in the skin prefigure striped colouration of zebrafish. Nat Cell Biol 16:607–614.

Sisley K, Curtis D, Rennie IG, Rees RC. 1993. Loss of heterozygosity of the thyroid hormone receptor B in posterior uveal melanoma. Melanoma Res 3:457–461.

Sosa KC, Colanesi S, Muller J, Schulte-Merker S, Stemple D, Elizabeth Patton E, Kelsh RN. 2018. Endothelin receptor Aa regulates proliferation and differentiation of Erb-dependant pigment progenitors in zebrafish. bioRxiv. doi:10.1101/308221

Spiewak JE, Bain EJ, Liu J, Kou K, Sturiale SL, Patterson LB, Diba P, Eisen JS, Braasch I, Ganz J, Parichy DM. 2018. Evolution of Endothelin signaling and diversification of adult pigment pattern in Danio fishes. PLoS Genet 14:e1007538.

Suster ML, Abe G, Schouw A, Kawakami K. 2011. Transposon-mediated BAC transgenesis in zebrafish. Nat Protoc 6:1998–2021.

Thisse C, Thisse B, Postlethwait JH. 1995. Expression of snail2, a second member of the zebrafish snail family, in cephalic mesendoderm and presumptive neural crest of wild-type and spadetail mutant embryos. Dev Biol 172:86–99.

Toews DPL, Hofmeister NR, Taylor SA. 2017. The Evolution and Genetics of Carotenoid Processing in Animals. Trends Genet 33:171–182.

Toomey MB, Lopes RJ, Araújo PM, Johnson JD, Gazda MA, Afonso S, Mota PG, Koch RE, Hill GE, Corbo JC, Carneiro M. 2017. High-density lipoprotein receptor SCARB1 is required for carotenoid coloration in birds. Proc Natl Acad Sci U S A 114:5219–5224.

Toomey MB, McGraw KJ. 2007. Modified saponification and HPLC methods for analyzing carotenoids from the retina of quail: implications for its use as a nonprimate model species. Invest Ophthalmol Vis Sci 48:3976–3982.

Trapnell C, Cacchiarelli D, Grimsby J, Pokharel P, Li S, Morse M, Lennon NJ, Livak KJ, Mikkelsen TS, Rinn JL. 2014. The dynamics and regulators of cell fate decisions are revealed by pseudotemporal ordering of single cells. Nat Biotechnol 32:381–386.

Usui Y, Kondo S, Watanabe M. 2018. Melanophore multinucleation pathways in zebrafish. Dev Growth Differ 60:454–459.

Van de Putte T, Maruhashi M, Francis A, Nelles L, Kondoh H, Huylebroeck D, Higashi Y. 2003. Mice lacking ZFHX1B, the gene that codes for Smad-interacting protein-1, reveal a role for multiple neural crest cell defects in the etiology of Hirschsprung disease-mental retardation syndrome. Am J Hum Genet 72:465–470.

Watanabe M, Iwashita M, Ishii M, Kurachi Y, Kawakami A, Kondo S, Okada N. 2006. Spot pattern of leopard Danio is caused by mutation in the zebrafish connexin41.8 gene. EMBO Rep 7:893–897.

Watanabe M, Kondo S. 2015. Is pigment patterning in fish skin determined by the Turing mechanism? Trends Genet 31:88–96.

Williams JS, Hsu JY, Rossi CC, Artinger KB. 2018. Requirement of zebrafish pcdh10a and pcdh10b in melanocyte precursor migration. Dev Biol. doi:10.1016/j.ydbio.2018.03.022

Zhang YM, Zimmer MA, Guardia T, Callahan SJ, Mondal C, Di Martino J, Takagi T, Fennell M, Garippa R, Campbell NR, Bravo-Cordero JJ, White RM. 2018. Distant Insulin Signaling Regulates Vertebrate Pigmentation through the Sheddase Bace2. Dev Cell 45:580–594.e.

Ziegler I. 2003. The pteridine pathway in zebrafish: regulation and specification during the determination of neural crest cell-fate. Pigment Cell Res 16:172–182.

Ziller C, Dupin E, Brazeau P, Paulin D, Le Douarin NM. 1983. Early segregation of a neuronal precursor cell line in the neural crest as revealed by culture in a chemically defined medium. Cell 32:627–638.

